# A normalization strategy for extreme proteome regulation reveals distinct targets of eIF4A1/2 and their inhibitor silvestrol

**DOI:** 10.64898/2026.11.16.767045

**Authors:** Toman Borteçen, Luca Gaessler, Robert Kalis, Elena Markeviciute, Johannes Zuber, Azmal Syed Ali, Jeroen Krijgsveld

## Abstract

Cells rapidly adjust protein synthesis in response to stress, signalling and drugs, often converging on the cell’s translation machinery affecting the synthesis of multiple proteins simultaneously. Nascent proteome analysis, combining pulsed SILAC labelling with enrichment of newly synthesized proteins (NSPs), can measure these changes proteome-wide. However, quantitative proteomics relies on normalization methods that assume most proteins remain unchanged. When protein synthesis shifts globally, this assumption fails: normalization then erases the biological effect, whereas omitting it leads to misinterpretation of data. Since no approach has so far reliably preserved global changes in protein synthesis, we developed labelled protein ratio (LPR) normalization, which uses the ratio of newly synthesized to pre-existing protein in the global proteome as an internal scaling factor. Two SILAC benchmarks showed that this factor is accurate down to a few percent of labelled protein and recovers defined global differences, whereas twelve established normalization methods did not. Applied to cycloheximide treatment, LPR normalization revealed a 57% decrease in synthesis, which median normalization reported as balanced up- and down-regulation with spurious increases in multiple proteins. After depletion of the core translation initiation factor eIF4A1, LPR normalization uncovered a broad translational repression that median normalization largely masked, while eIF4A inhibition with silvestrol showed a divergent pattern, suggesting secondary effects. LPR normalization thus provides a general strategy for quantifying global changes in protein synthesis.

## Introduction

Proteome analysis is a powerful approach to examine changes in protein abundance to gain a systems understanding how cells respond to specific perturbations. Various experimental strategies have been devised to perform protein quantitation in a proteome-wide manner, either based on stable isotope labelling or using label-free approaches, in conjunction with computational tools to appropriately analyse such data. The main tasks of such tools include data normalization across conditions, determination of fold-changes in protein abundance, and assigning a statistical measure for each protein to indicate the significance of this change. This general strategy has been successfully used in countless studies; however, problems arise in scenarios that violate key assumptions, namely that the majority of proteins remain unchanged, and that changes in protein expression follow a standard normal distribution (1–3). These criteria are met in most biological settings where proteome responses are relatively mild. However, instances of extreme proteome regulation may produce a complete shift in abundance distribution. This may occur in processes that strongly depend on protein synthesis, or upon sharp induction of protein degradation or shutdown of protein translation as a response to stress (starvation, ER stress). In each of these scenarios, conventional (median) normalization fails, and to the best of our knowledge, no suitable approach has yet been developed to handle such proteomic data in a statistically sound manner.

Analysis of newly synthesized proteins (NSPs) via nascent proteome analysis enables direct assessment of perturbation-induced changes in protein synthesis. A strong asset of NSP analysis is that proteome alterations become apparent as an immediate consequence of the perturbation, before this can be detected at the level of global protein expression (4). Methods for the quantitative analysis of NSPs rely on metabolic labelling with either stable isotope-containing amino acids (5), non-canonical amino acids (6), or combinations thereof (7), enabling NSPs to be distinguished from pre-existing proteins. Combining pulsed SILAC with incorporation of the methionine analogue azidohomoalanine (AHA) is particularly powerful as it provides an azide handle for click-chemistry enrichment of NSPs, improving the sensitivity to quantify changes in protein synthesis proteome-wide (4, 7). We recently developed an automated implementation of this strategy to facilitate sample handling and enhance throughput of NSP enrichment, and used plexDIA (8) to achieve increased proteome coverage and quantitative accuracy in single shot analyses (9).

Despite these advances, normalization of nascent proteome data poses an additional challenge that has to our best knowledge not been addressed yet. Most normalization methods for quantitative proteomics require that the abundance of most proteins remains unchanged between the tested conditions (2, 10, 11). However, this assumption is violated when protein synthesis is globally reduced or increased, as frequently occurs when cells adapt to adverse conditions. For instance, cellular stress conditions such as oxidative stress (12), nutrient starvation (13), cold (14) and heat shock (15) lead to the global inhibition of protein synthesis. This mainly occurs at the level of translation initiation via phosphorylation of eIF2α and inhibition of the 5’-cap-binding protein complex eIF4F (16), but can also involve pausing of elongation (17). In addition, mTOR, the integrator of a variety of cellular pathways, controls translation of a wide range of downstream targets by phosphorylating 4E-BP1 to regulate eIF4E (18). Prolonged stress can trigger irreversible translational arrest through targeted cleavage of translation initiation factors by apoptotic proteases (19). Protein synthesis is also cell cycle-regulated, being globally inhibited during interphase (20) and is further reduced when cells enter quiescence in G0 – an important mechanism by which cells prevent premature exhaustion or develop drug resistance (21). The translational machinery is central to various pathologies, most notably during viral infection: infected cells shut down translation to prevent viral replication, while viruses have evolved a range of mechanisms to hijack the very same machinery to translate their own transcripts (22). Finally, numerous toxins produced by fungi, plants and other organisms are natural protein-synthesis inhibitors; they are the origin of several major classes of antibiotics (e.g. tetracyclines, macrolides) (23), or serve as tool compounds in biochemical experiments (e.g. cycloheximide) to globally inhibit translation in eukaryotic cells to investigate protein turnover or translational regulation (24). Studying any of these processes and biological systems at the proteome level requires handling large and biologically meaningful shifts in the abundance of newly synthesized proteins, which regular median normalization would erase (10, 11).

Translation initiation is tightly regulated by several dozen eukaryotic translation initiation factors (eIFs) that recognize the mRNA cap, facilitate mRNA unwinding, and recruit the ribosome for protein synthesis (25). Translation is a core process for any cell, however is of key importance for cancer cells as they depend on it to maintain cell growth and proliferation. Indeed, various eIFs have been found to be deregulated across diverse cancer types (26, 27), making them attractive therapeutic targets to inhibit tumor growth (28). An eIF that has gained particular interest is eIF4A, a member of the 5’-cap-binding eIF4F complex that controls global protein synthesis by unwinding secondary mRNA structures via its RNA helicase activity. Its two isoforms (eIF4A1 and eIF4A2) share 90% sequence homology in humans and are used interchangeably in the eIF4F complex, however only eIF4A1 is essential, and they target different transcripts with divergent functions (29–33). Silvestrol is a member of the Rocaglate-family of compounds that inhibits translation by clamping eIF4A on mRNA, preventing the scanning of the 43S pre-initiation complex (PIC) or trapping the eIF4F complex at the 5’-cap (34, 35). In cell culture models and *in vivo* experiments, silvestrol inhibits protein synthesis and tumor growth (36, 37). At the same time, proteomic studies have indicated that silvestrol has broader implications in proteome adaptation than only inhibiting EIF4A, since proteome effects after silvestrol treatment could not be replicated by eIF4A depletion (38).

Since rapid and profound proteome remodelling is a key mechanism of cells to adapt to adverse conditions, and since proteomics has the unique ability to directly quantify these changes on a proteome-wide scale, there is a clear need for robust analytical approaches. However, current data-handling strategies are poorly suited to proteomic measurements under conditions of strong translational perturbation. Therefore, we here introduce a normalization approach termed labelled protein ratio (LPR) normalization, which enables proteome-wide quantification of changes in protein synthesis by combining nascent- and global proteome analysis. We apply this approach to quantify shutdown of protein translation by cycloheximide, and to compare changes in protein synthesis induced by depletion of eIF4A1/2 or by silvestrol treatment. LPR normalization and differential proteome analysis revealed substantial differences in protein synthesis between eIF4A1 and eIF4A2 depletion and identified a distinct signature for silvestrol treatment, underscoring that these perturbations are not mechanistically equivalent. Together, these data show that LPR normalization is an effective and broadly applicable strategy for studying cellular responses that involve profound translational remodelling.

## Results

### Labelled protein ratio normalization preserves global changes in protein synthesis

Protein synthesis is highly dynamic and can change globally within hours. Cellular stress, nutrient limitation, signalling through mTOR, viral infection, or translation inhibitors can repress or activate the synthesis of many proteins simultaneously, resulting in a global shift as schematically indicated in **Figure 1A**. Standard median normalization then re-centres the distribution at zero and removes the biological shift (**Figure 1A**). As a consequence, proteins whose synthesis is reduced are reported as unchanged, and proteins that are reduced less than average appear up-regulated. To avoid such artifacts, normalization could be omitted in principle, however this requires mixing of exactly the same amounts of protein from control and treated cells before NSP enrichment. In practice this is difficult to achieve, particularly with low-input samples where small errors in protein quantification or pipetting translate directly into a systematic offset of all ratios.

**Figure 1.**
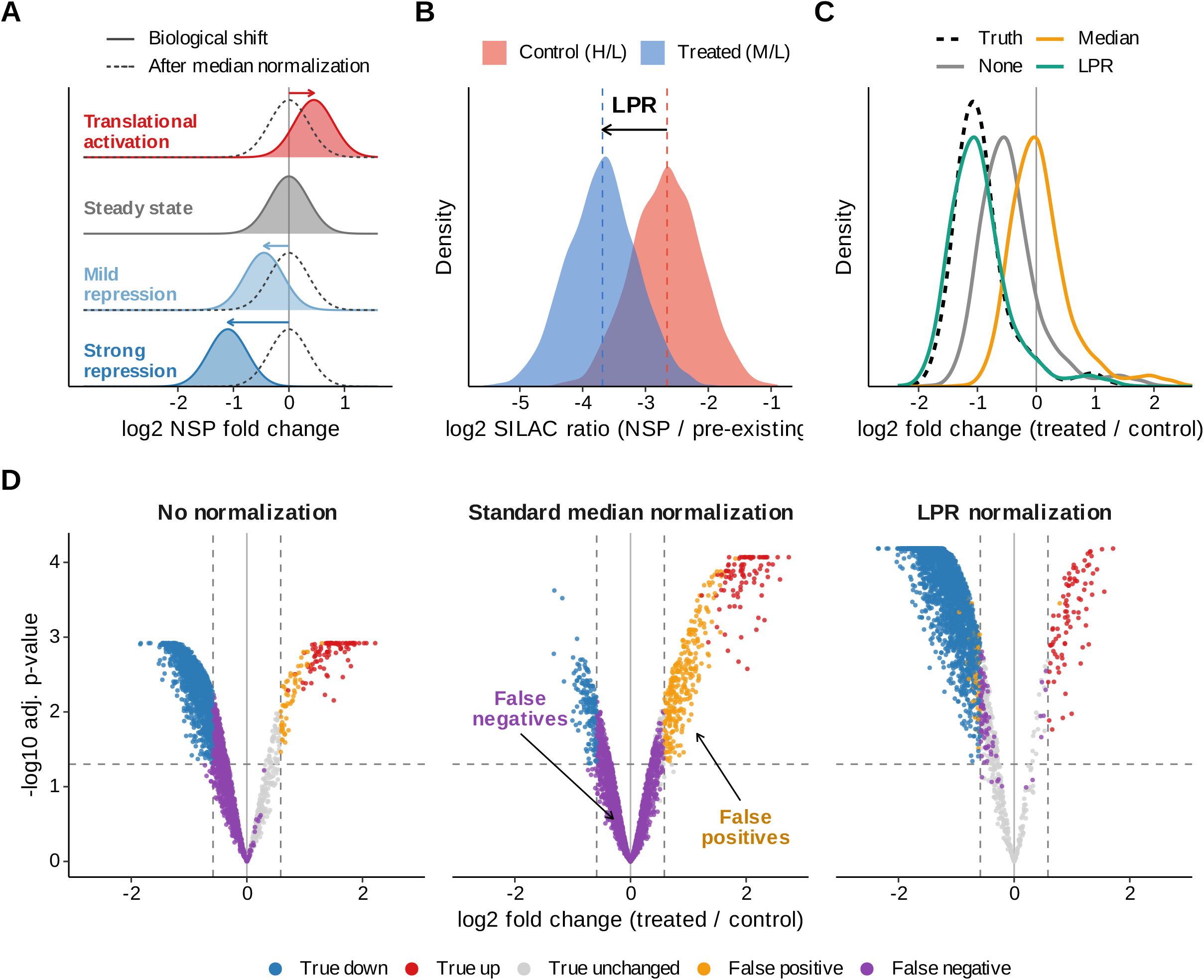
Principle of LPR normalization and comparison with standard normalization strategies. A) Global changes in protein synthesis (activation, steady state, mild or strong repression) shift the distribution of NSP log2 fold changes (filled). Standard median normalization re-centres each distribution at zero (dashed lines), thus removing the biological effect. B) Determination of the LPR factor. Control and treated cells are pulse-labelled with heavy (H) and Medium (M) SILAC amino acids, and pre-existing proteins remain light (L). Before NSP enrichment, an aliquot of each lysate is analysed as a global proteome. The difference between the median log2 M/L and H/L ratios (dashed lines, arrow) reports the global change in synthesis and is used as the LPR factor. C) Distributions of NSP log2 fold changes (treated/control) for the ground truth (dashed black) and without normalization (grey), or after standard median normalization (orange) or LPR normalization (teal). D) Volcano plots for the three strategies, with proteins coloured by agreement with the ground truth: true down (blue), true up (red), true unchanged (grey), false positive (orange) and false negative (purple). Dashed lines indicate log2 fold change = 0.58 and adjusted p = 0.05. Data in B–D are simulated (see Methods).

To retain true global changes while correcting for such technical variation, we developed labelled protein ratio (LPR) normalization. The approach combines nascent and global proteome analysis of the same samples: control and treated cells are pulse-labelled with different SILAC amino acids, here heavy (H) for control and Medium (M) for treated cells, so that NSPs carry the label while pre-existing proteins in either condition remain light (L). Before NSP enrichment, an aliquot of each lysate is analysed separately as a global proteome. In these samples, the ratio of labelled to unlabelled protein (H/L for control, M/L for treated) reflects how much protein was synthesized during the pulse relative to the pre-existing proteome (**Figure 1B**). Because the light, pre-existing proteome serves as a common internal reference, the difference between the median log2 M/L and the median log2 H/L ratio directly reports the global change in protein synthesis. We define this difference, which corresponds to log2 M/H, as the LPR factor. The remaining lysates are then mixed, NSPs are enriched and subjected to proteome analysis. Median-centring of the NSP ratios (M/H) in the resulting data removes the mixing error, but also the global change in protein synthesis, which is then restored by shifting the fold-change distribution by the LPR factor.

To illustrate how different normalization strategies affect the outcome, we simulated a nascent proteome experiment with a known ground truth, in which a treatment globally represses protein synthesis, and where control and treated lysates are mixed in slightly unequal amounts before NSP enrichment (**Figure 1C, D; Methods**). Without normalization, the mixing error offset the entire fold-change distribution towards zero, so that many truly repressed proteins were missed (**Figure 1D, left panel**). Standard median normalization corrected the mixing error but also removed the biological effect. As a result, most repressed proteins were no longer recognized as such (false negatives), and proteins that were reduced less than average appeared up-regulated (false positives) (**Figure 1D, middle panel**). In contrast, LPR normalization restored the true fold-change distribution and identified repressed and induced proteins with few false calls (**Figure 1D, right panel**). The LPR factor estimated from the simulated global proteome (log2 −1.03) closely matched the true global change in protein synthesis implemented in the simulation (median log2 fold change −1.02).

### Benchmarking accuracy and dynamic range of LPR normalization

To derive the scaling factor for LPR normalization, H/L and M/L ratios must be quantified accurately and linearly. This is a demanding requirement in pSILAC samples that contain only a small fraction of newly synthesized proteins (NSPs), as occurs when protein synthesis is strongly repressed. Therefore, we benchmarked quantification accuracy and dynamic range of quantification using a dilution series of samples with different fractions of heavy-, medium- and light labelled proteins with mixing ratios ranging from equal proportions to a 94-fold difference (**Figure 2A; Supplementary Table 1**). The samples were analysed via a 90 min DIA method (70 min active gradient) on a Q Exactive HF Orbitrap mass spectrometer, using the plexDIA module of DIA-NN for data analysis.

**Figure 2.**
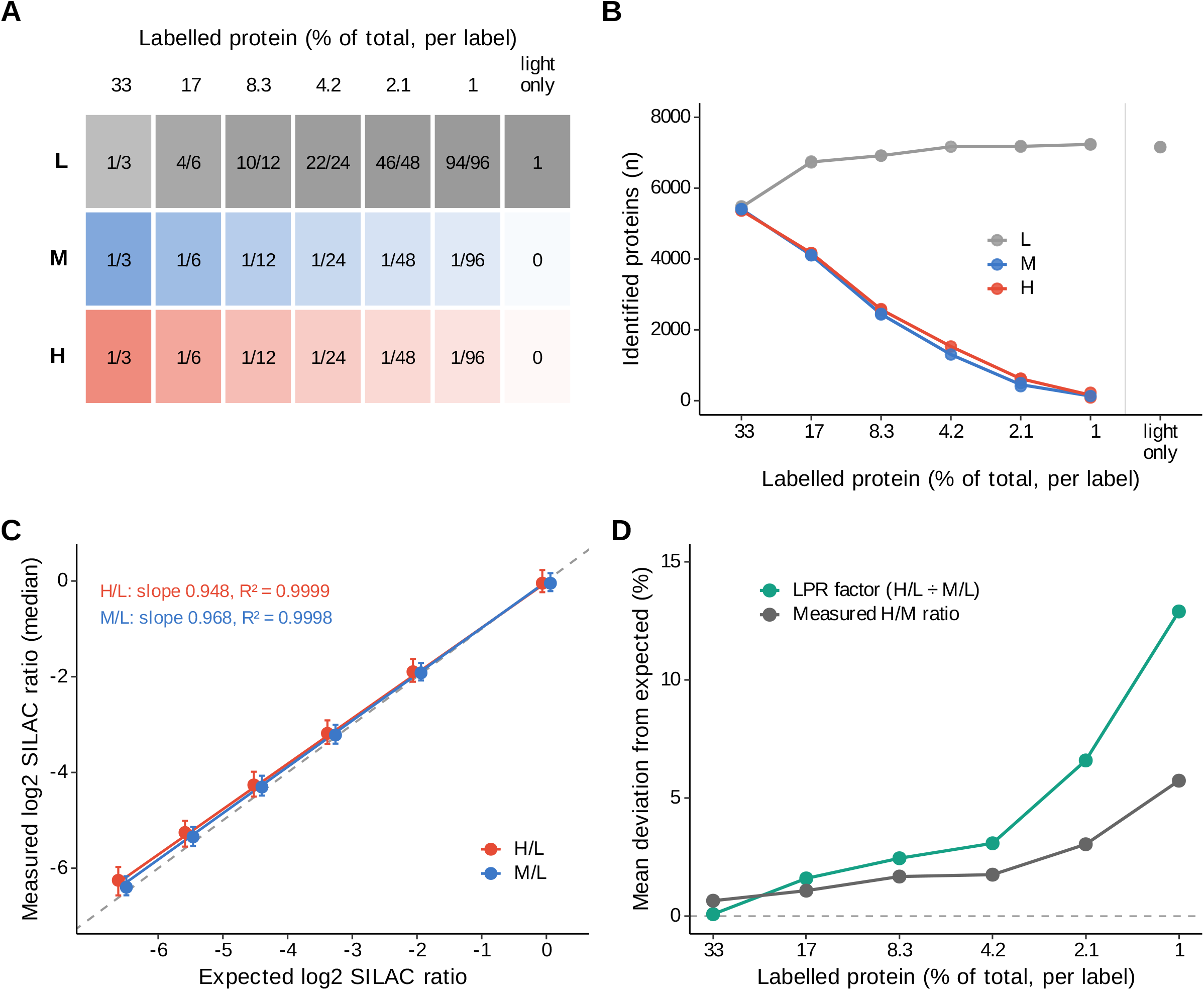
Accuracy of SILAC ratio quantification and of the LPR factor across a dilution series of labelled protein. A) Composition of the benchmark samples. Light (L), medium (M) and heavy (H) SILAC-labelled HeLa lysates were mixed so that each labelled channel made up 33% to 1% of the total protein, plus a light-only sample. Numbers indicate the fraction of each label. Samples were analysed by 90 min DIA on a Q Exactive HF and processed with DIA-NN (plexDIA); n = 2 replicates per sample. B) Number of proteins identified in each SILAC channel; points show individual replicates and lines the mean. C) Measured versus expected median log2 H/L and M/L ratios, shown as the mean of the replicate medians. Error bars indicate the interquartile range of protein ratios (mean of replicates), solid lines show linear fits (slopes and R² indicated), and the dashed line indicates perfect agreement. D) Deviation of the LPR factor (median log2 H/L minus median log2 M/L) and of the directly measured median H/M ratio from the expected 1:1 ratio, shown as mean deviation (%) across replicates per sample; at 1% labelled protein, LPR factor and H/M ratio are based on one replicate.

Due to the use of stringent channel- and translated q-value filters (< 0.01) and lack of re-quantify in DIA-NN, the number of identified SILAC-labelled proteins correlates with the abundance of proteins with the respective label in the sample, as we previously described (9). Around 7000 proteins were identified in the sample that exclusively consisted of unlabelled (light) proteins (**Figure 2B**). This number decreased with increased proportions of medium and heavy-labelled proteins, as expected, with approximately 5500 proteins identified for each labelling state in the 1:1:1 mixture. The number of detected labelled proteins decreased as their proportion dropped from 1/3 (33%) to 1/96 (1%) in the dilution series, again as expected (red and blue in **Figure 2B**).

More importantly, the median SILAC ratios showed very high linearity (R2 > 0.999) for both heavy and medium SILAC channels (**Figure 2C**), with high quantitative accuracy across the intensity range even when labelled proteins represented only 1% of the total sample abundance (**Supplementary Figure 1**; per-protein linearity in **Supplementary Figure 2A**). This is also reflected by the small CVs (median around 5%, **Supplementary Figure 2B**) and the observation that deviation from the expected protein ratio was small (median 0.2–0.4 log2 units), largely independent of the absolute value of the ratio (**Supplementary Figure 2C**). Applying the LPR normalization approach to the benchmark samples showed that the calculated median H/M SILAC ratios were highly accurate, deviating by 3% or less from the expected values down to 4.2% labelled protein (**Figure 2D**). However, the mean deviation increased in samples with very low fractions of labelled proteins, reaching 13% for the LPR factor and 6% for the measured H/M ratio at 1% labelled protein (single replicate) (**Figure 2D** and **Supplementary Figure 2C**). The suitable quantification range should therefore be carefully assessed for each instrument, acquisition method and software.

### Benchmarking of LPR normalization before and after NSP enrichment

Next, we set up an experiment to better reflect an experimental scenario where the fraction of labelled protein is unknown, but that still spans a wide range. In addition, we aimed to evaluate the performance of LPR normalization for low H/L and M/L ratios before and after enrichment of NSP. Therefore, we pulse-labelled HeLa cells for 4 h using AHA together with either heavy or medium lysine and arginine-containing medium and mixed the lysates at defined H:M ratios (**Figure 3A; Supplementary Table 2**). In subsequent global proteome analysis (i.e. prior to NSP enrichment), the number of identified L, M, and H-labelled proteins correlated well with their relative abundance in the samples (**Figure 3B, left panel**). In the 10:90 and 90:10 H:M mixtures, no H/M ratios could be quantified (**Figure 3C, left panel**), because the H- or M-labelled proteins, respectively, were not identified at this very low abundance (**Figure 3B, left panel**).

**Figure 3.**
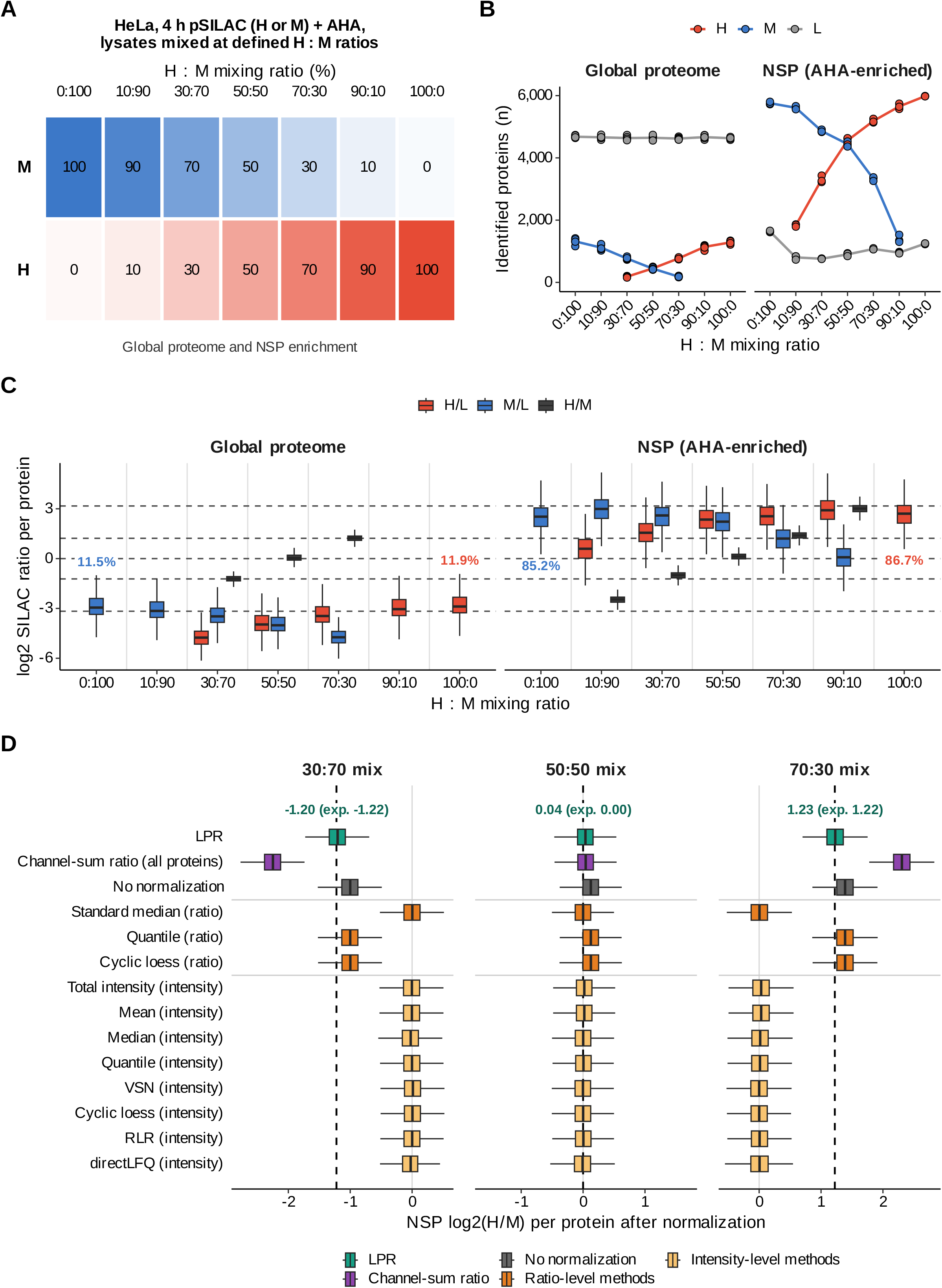
Recovery of defined global differences in labelled protein by LPR normalization in a pulsed SILAC/AHA mixing benchmark. A) HeLa cells were pulse-labelled for 4 h with either heavy (H) or medium (M) SILAC amino acids together with AHA, and the lysates were mixed at defined H:M ratios (numbers indicate % of each). Mixed samples were analysed as global proteome (n = 6) and after NSP enrichment (n = 3) by 130 min DIA method (110 min active gradient) on an Orbitrap Fusion. B) Number of proteins quantified in the light (L), M and H channels in the global proteome and in the NSP-enriched samples; points show individual replicates and lines the mean. C) Log2 SILAC ratios per protein (H/L, M/L and H/M; mean of replicates) in the global proteome and NSP samples for each mixing ratio. Dashed lines indicate the expected log2 H/M ratios of the mixes (0, ±1.22 and ±3.17). Numbers indicate the fraction of labelled protein in the unmixed samples, calculated from the median ratio (m) as 2^m / (1 + 2^m). Ratios are shown only where at least 20 proteins were quantified. D) NSP log2 H/M ratios per protein after LPR normalization, after channel-sum ratio normalization, without normalization, and after ratio-level and intensity-level normalization methods, for the 30:70, 50:50 and 70:30 mixes. Dashed lines indicate the expected log2 H/M ratio. Numbers show the median after LPR normalization and the expected value. Boxes show the interquartile range with the median, and whiskers 1.5× the interquartile range (C, D).

Indeed, based on the median H/L and M/L ratios of the unmixed samples (100:0 and 0:100), labelled NSP accounted for 11–12% of the total protein in the non-enriched samples after the 4 h pulse (**Figure 3C, left panel**), so that the minor label in the 10:90 and 90:10 mixtures would be expected to account for only about 1.2% of total protein. This is similar to the lowest fraction tested in the dilution benchmark, albeit on a different instrument, at which only 100 to 200 proteins could still be quantified (**Figure 2B**). In contrast, after enrichment of NSPs from the same mixed samples, the number of identified M- and H-labelled proteins was notably higher than in the global proteome prior to enrichment, while the opposite was true for light proteins (**Figure 3B, right panel**), reflecting effective enrichment of labelled NSPs and depletion of pre-existing proteins. Accordingly, the fraction of H- and M-labelled protein increased to 87% and 85%, respectively, after NSP enrichment (**Figure 3C, right panel**). Notably, both in the global and NSP proteome, no M- or H-labelled proteins were identified in samples that did not contain that label in the mixture (**Figure 3B**), indicating high specificity of the analysis.

Applying LPR normalization based on the ratio of the median H/L and M/L SILAC ratios in non-enriched samples shifted the median-centred NSP H/M ratios to log2 values of −1.20, 0.04 and 1.23 for the 30:70, 50:50 and 70:30 mixtures, closely matching the expected values of −1.22, 0 and 1.22 (**Figure 3D**). Importantly, regular median normalization as well as eleven other normalization methods failed in this task, most of them centring the distribution around 0 irrespective of the mixing ratio, while the channel-sum ratio approximately doubled the expected values (**Figure 3D**). Ratio-based quantile and cyclic loess normalization left the non-normalized ratios unchanged and therefore retained their offset from the expected values (−1.00 and 1.39), which LPR normalization corrected (**Figure 3D**). This shows that a wide range of ratio- and intensity-based normalization methods struggle to handle imbalanced NSP data.

These results establish that plexDIA can accurately quantify median SILAC ratios to enable LPR normalization across challenging samples, including those with strongly skewed protein distributions or low fractions of labelled protein. This indicates that the approach should be applicable to scenarios using short pSILAC labelling and strong inhibition of protein synthesis.

### Quantifying global inhibition of mRNA translation in response to cycloheximide treatment

To test LPR normalization in a cellular perturbation experiment, we treated HeLa cells with CHX to globally inhibit protein synthesis. CHX is a potent translation inhibitor, however a 4 h treatment with a high dose of CHX (100 µg/mL) did not lead to a noticeable change in protein abundance when investigated by global label-free proteomics (none of 6,282 quantified proteins changed significantly, **Supplementary Figure 3A**). To investigate inhibition of protein synthesis in more detail, we treated cells with different doses of CHX (from 0.1 to 100 µg/mL) for 4 h, with simultaneous pulsed SILAC and AHA labelling, followed by click-based enrichment of NSPs and LC-MS analysis (**Figure 4A**).

**Figure 4.**
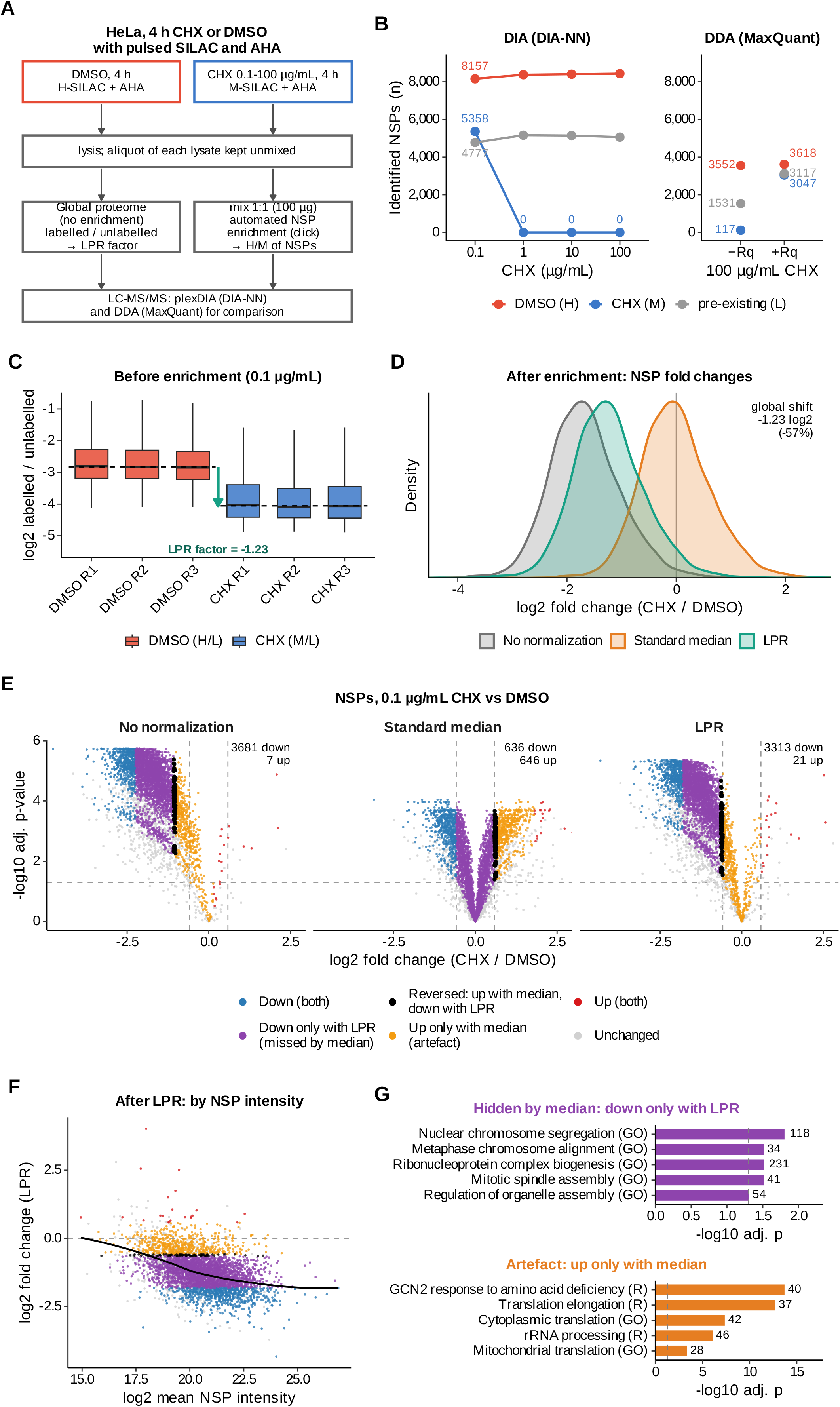
LPR normalization reveals global inhibition of protein synthesis by cycloheximide. A) HeLa cells were treated with DMSO or cycloheximide (CHX, 0.1–100 µg/mL) for 4 h, and NSPs were labelled with heavy (H; DMSO) or medium (M; CHX) SILAC amino acids together with AHA. An aliquot of each lysate was analysed as a global proteome to determine the LPR factor. The remaining lysates were mixed 1:1 (100 µg total protein) for NSP enrichment, and all samples were analysed by plexDIA (DIA-NN); n = 3 replicates per condition. For comparison, the 100 µg/mL CHX sample was also analysed by DDA. B) Number of NSPs identified in the H (DMSO), M (CHX) and light (L, pre-existing) channels after enrichment, at each CHX concentration (left, DIA; mean of replicates). Right, DDA analysis of the 100 µg/mL CHX sample, processed with MaxQuant without (−Rq) or with (+Rq) the re-quantify option. C) Log2 ratios of labelled over unlabelled protein in the global proteome of DMSO-(H/L) and CHX-treated (0.1 µg/mL; M/L) cells, per replicate. Dashed lines indicate the mean of the replicate medians per condition, and the arrow the resulting LPR factor. D) Distribution of NSP log2 fold changes (CHX/DMSO) without normalization, after standard median normalization and after LPR normalization. E) Volcano plots of NSPs without normalization, after standard median normalization and after LPR normalization. Proteins are coloured by their regulation with standard median and LPR normalization (|log2 fold change| > 0.585, adjusted p < 0.05, CV < 20%); numbers indicate down- and up-regulated NSPs. F) NSP log2 fold changes after LPR normalization plotted against mean NSP intensity; the black line shows a loess fit. Colours as in E. G) GO Biological Process (GO) and Reactome (R) terms enriched among NSPs down-regulated only with LPR (hidden by median normalization) and among NSPs up-regulated only with median normalization (artefact). Numbers indicate the NSPs per term, and dashed lines indicate an adjusted p-value of 0.05 (one-sided Fisher’s exact test, Benjamini-Hochberg correction; background: all quantified NSPs).

Strikingly, not a single NSP from CHX-treated cells was identified at 1–100 µg/mL CHX (**Figure 4B, left**), indicating that indeed protein translation was shut down completely. At the lowest CHX concentration (0.1 µg/mL), >5,000 NSPs were identified, compared to >8,000 in the DMSO-treated controls (**Figure 4B, left**), indicating that 0.1 µg/mL CHX partially inhibits protein synthesis. In stark contrast to the absence of NSPs using DIA-based analysis, a DDA-based strategy identified >100 NSPs in HeLa cells treated with 100 µg/mL CHX (**Figure 4B, right**). Using the re-quantify function, which assigns a quantity to the missing SILAC partner from background signal, this number even increased to >3,000, i.e. approaching the number of NSP identifications in DMSO-treated cells analysed by DDA (**Figure 4B, right**). These data highlight the need for caution when processing and interpreting DDA data in this context, while demonstrating the power of DIA-NN in handling triple-SILAC data.

SILAC ratios of NSP over pre-existing proteins prior to NSP enrichment were smaller in cells treated with 0.1 µg/mL CHX than in DMSO controls (**Figure 4C**), as expected because of inhibited translation. The difference between these ratios was used as an LPR scaling factor for the normalization of the click-based enriched nascent proteome data. Specifically, the median log2 ratios were −4.05 and −2.83 in CHX- and DMSO-treated samples, respectively (**Figure 4C**), resulting in a log2 scaling factor of −1.23. Shifting the distribution of SILAC ratios in the enriched NSP sample by this factor (**Figure 4D**) leads to a volcano plot indicating strong and significant downregulation of the large majority of proteins upon CHX treatment (3,313 of 4,539 NSPs; **Figure 4E; Supplementary Table 3**). Without normalization, repression was overestimated (median log2 fold change −1.70 vs −1.27 after LPR normalization), reflecting unequal input of the two SILAC channels, whereas standard median normalization removed the effect entirely, leaving similar numbers of up- and down-regulated proteins (646 and 636) (**Figure 4D, E**). Interestingly, NSPs with higher intensity appeared to be more strongly downregulated than those with lower intensity (**Figure 4F**). Not surprisingly, the subset of 1,134 proteins that were identified in proteome samples without click chemistry-based enrichment and that were subsequently used for the determination of the scaling factor were highly abundant in HeLa cells (**Supplementary Figure 3B**). Importantly, this subset of proteins did not differ in their overall abundance between the two conditions (median log2 fold change 0.01), thus validating the main assumption of the normalization strategy (**Supplementary Figure 3B**). Functionally, NSPs whose repression was only detected after LPR normalization were enriched for chromosome segregation, mitotic spindle assembly and ribonucleoprotein complex biogenesis, whereas proteins that appeared up-regulated only after median normalization were enriched for cytoplasmic and mitochondrial translation and the GCN2 response to amino acid deficiency (**Figure 4G**; all regulation groups and proteins with reversed direction in **Supplementary Figure 3C, D**).

These results demonstrate that LPR normalization accurately captures extreme and imbalanced proteome changes after inhibition of global protein synthesis, correcting technical bias without removing the biological effect. Moreover, and in contrast with DDA analysis, loss of protein synthesis can be accurately measured through SILAC plexDIA.

### Nascent proteome analysis upon eIF4A1/2 depletion and inhibition by silvestrol

Having established a strategy to detect and quantify inhibition of protein synthesis on a proteome-wide scale, we aimed to apply it to investigate how eIF4A1 and eIF4A2 shape the nascent proteome. These eIFs are paralogous RNA helicases that participate in the cap-binding eIF4F complex in a mutually exclusive manner during translation initiation. Since eIF4A1 (but not eIF4A2) is essential for cell survival (29, 39), we established an inducible CRISPR-Cas9 knock-out (KO) system in RKO cells allowing depletion of either protein (see Methods). In a complementary approach, we inhibited eIF4A activity with silvestrol.

To ensure comparable experimental conditions between eIF4A depletion and inhibition, we assessed cell viability in RKO cells upon induced KO of eIF4A1 and different doses of silvestrol over a time course of 72 h (**Supplementary Figure 4**). As a condition that did not induce major cell death, we chose to treat RKO cells for 48 h either with doxycycline (dox) to induce KO of eIF4A1/2, or with 3 nM silvestrol, which is in line with previous studies (37, 40). These samples were then subjected to nascent proteome analysis by shifting cells to AHA- and SILAC amino acid-containing media during the last 6 h of dox/silvestrol treatment, followed by NSP enrichment and LC-MS/MS. In addition, an aliquot of each sample was analysed prior to enrichment to determine the LPR normalization factor accounting for global changes in protein synthesis. This latter experiment showed that depletion of eIF4A1 led to a decrease in overall NSP levels by on average 18%, while a minor increase (3%) was observed upon eIF4A2 depletion and a minor decrease (4%) upon silvestrol treatment (**Figure 5A**). When applying these as LPR normalization factors after NSP enrichment (**Figure 5B; Supplementary Table 4**), depletion of eIF4A1 showed a strong impact on the nascent proteome with 890 proteins being inhibited and 84 enhanced in translation, while a much milder response was observed for eIF4A2 (5 inhibited, 27 enhanced). Silvestrol treatment had an intermediate effect, with 296 proteins inhibited and 156 enhanced (**Figure 5C**). Of note, LPR normalization recovered many downregulated proteins not captured by median normalization, while eliminating spurious up-regulated proteins (for eIF4A1 depletion, 278 down and 271 up after median normalization) (**Figure 5C, D**). By extension, many functional categories represented by these proteins could be corrected (**Figure 5F**). The stark difference in proteome response between eIF4A1 and eIF4A2 depletion is in line with previous studies using proteomics and ribosome profiling (32, 38). Interestingly, depletion of eIF4A1 led to an increase of eIF4A2 in NSP- and overall protein abundance, while depletion of eIF4A2 did not change eIF4A1 (**Figure 5D, E, and Supplementary Figure 5**). This mirrors earlier observations both at the transcript and protein level (32, 38, 41), and confirms that LPR normalization did not skew this pattern.

**Figure 5.**
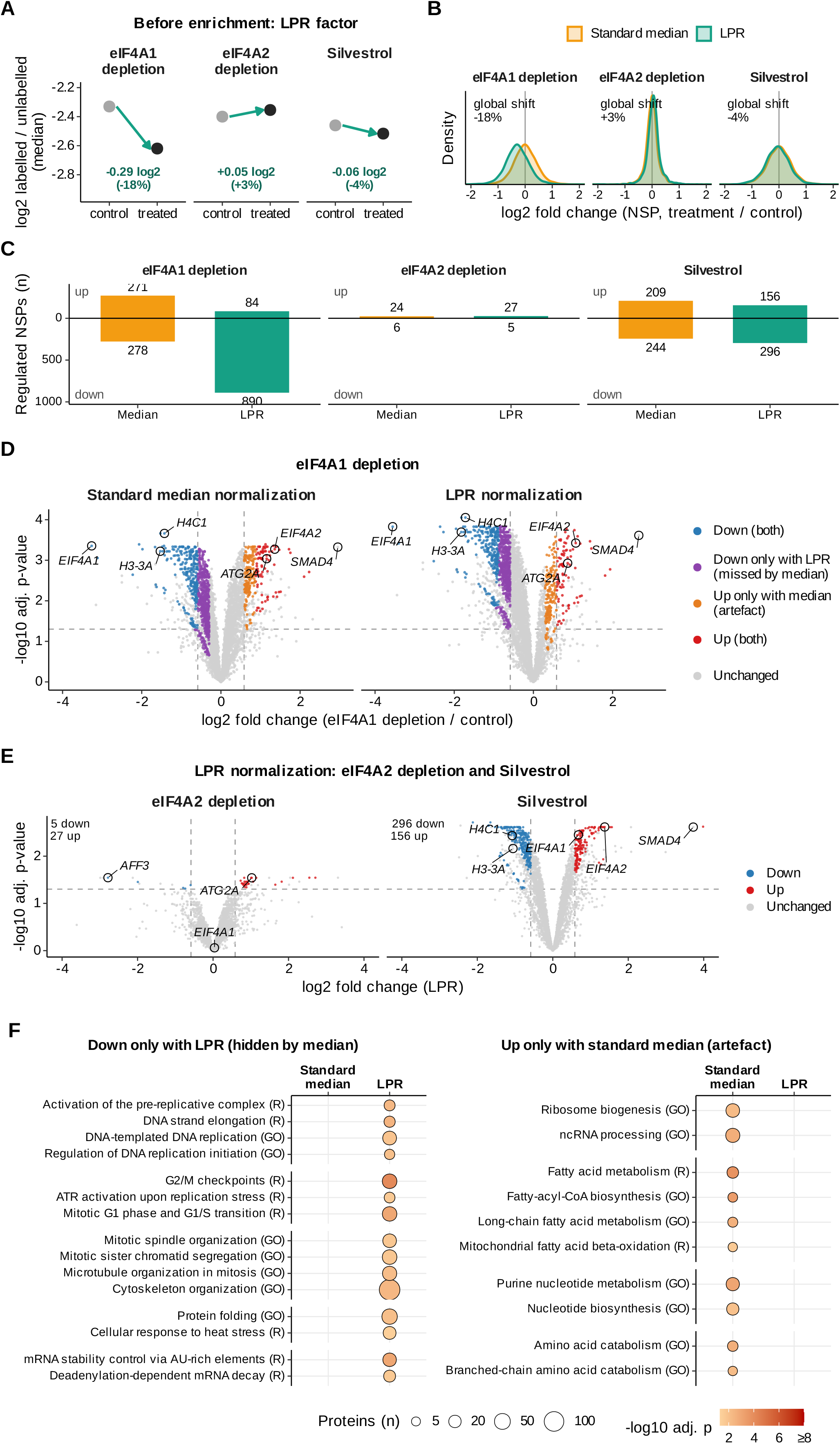
LPR normalization of nascent proteome data after eIF4A1 or eIF4A2 depletion and silvestrol treatment. **A)** LPR factors determined from the global proteome before NSP enrichment. Median log2 ratios of labelled over unlabelled protein are shown for control and treated RKO cells after 48 h of doxycycline-induced knockout of eIF4A1 or eIF4A2 (control: AAV1 knockout) or treatment with 3 nM silvestrol (control: DMSO). NSPs were labelled with SILAC amino acids and AHA during the last 6 h. Arrows and numbers indicate the LPR factor and the corresponding change in global protein synthesis. **B)** Distributions of NSP log2 fold changes (treated/control) after standard median normalization and after LPR normalization. **C)** Number of down- and up-regulated NSPs after standard median normalization and after LPR normalization. **D)** Volcano plots of NSPs after eIF4A1 depletion with standard median and LPR normalization. Proteins are coloured by their regulation with the two methods; selected proteins are labelled. **E)** Volcano plots of NSPs after eIF4A2 depletion and silvestrol treatment with LPR normalization. **F)** GO Biological Process (GO) and Reactome (R) terms enriched after eIF4A1 depletion among NSPs down-regulated only with LPR (left) and among NSPs up-regulated only with standard median normalization (right). Dot size indicates the number of proteins per term, and colour the adjusted p-value (one-sided Fisher’s exact test, Benjamini-Hochberg correction).

Silvestrol treatment induced up-as well as down-regulation of numerous proteins both in the global proteome (Supplementary Figure 5) and in the nascent proteome (Figure 5C, E), with only a 4% net decrease in NSPs (Figure 5A). This dichotomous behaviour to both induce or repress translation of mRNA subsets has been documented before in several studies using ribosome profiling (34, 42). Interestingly, both eIF4A1 and eIF4A2 were up-regulated upon silvestrol treatment (**Figure 5E and Supplementary Figure 5**). The balanced up- and down-regulation of proteins upon silvestrol treatment contrasts with the predominantly translational inhibition caused by eIF4A1 depletion. This challenges the notion that silvestrol is a specific eIF4A inhibitor, although several explanations may be possible (see Discussion). Pearson correlation between the nascent proteome changes of the three conditions was relatively low (R = −0.06 to 0.30; **Supplementary Figure 6A**), indicating complementary proteome effects between depletion of eIF4A1 or eIF4A2 and silvestrol treatment. Among the 890 proteins that were downregulated upon eIF4A1 depletion, only 108 were shared with silvestrol treatment. However, this represented the largest overlap among all comparisons, for both up- and down-regulated proteins, where some yielded only a few or even no proteins in common (**Supplementary Figure 6B**). For instance, histone proteins H3-3A and H4C1 were strongly decreased both in eIF4A1-depleted and silvestrol-treated cells, while the signalling protein SMAD4 was strongly increased (**Figure 5D, E**). Among the downregulated proteins upon eIF4A1 and eIF4A2 depletion only two overlapped, TLE5 and PRPF4B, which were also downregulated upon silvestrol treatment, while the autophagy-associated protein ATG2A was the only commonly upregulated protein (**Figure 5D, E; Supplementary Figure 6B**).

Collectively, these results indicate that LPR normalization effectively handles nascent proteomics data with large and modest systematic shifts in protein synthesis upon eIF4A1 and eIF4A2 depletion, respectively, and thereby avoids the artefactual regulation patterns produced by conventional median normalization. Moreover, the limited overlap between the proteins regulated after depletion of either paralog and those after silvestrol treatment underscores the complexity of translational control and of silvestrol’s mechanism of action.

## Discussion

Data normalization is central to any proteomic comparison of samples. Here we present LPR normalization, a strategy that quantifies proteome-wide changes accurately even when global protein synthesis is strongly altered. LPR combines nascent- and global proteome analysis in a SILAC plexDIA workflow to measure ratios of stable-isotope-labelled proteins. We applied it to proteome-wide changes induced by CXH-mediated global inhibition of protein synthesis, by depletion of the translation initiation factors eIF4A1 and eIF4A2, and by silvestrol-mediated inhibition of eIF4A. LPR corrects the skewed protein distributions where regular median normalization fails, and reveals distinct patterns of proteome regulation by eIF4A1, eIF4A2 and silvestrol.

Normalization methods for label-free and label-based proteomics assume that most proteins do not change between conditions (2, 10). This assumption fails when global protein synthesis is altered, which occurs in a wide range of stress conditions eliciting adaptation mechanisms that often converge on the translational machinery (16). Being able to quantify such changes reliably is important, since the processes involved (e.g. viral response, entry into quiescence, development of drug resistance) often are of (patho)physiological importance. Therefore, reliable protein quantification is crucial to mechanistically understand how cells adapt, or to design intervention strategies. In addition, skewed proteome distributions and imperfect normalization may be a more common phenomenon: conventional median normalization failed to resolve differences between induced pluripotent and embryonic stem cells because of disparities in protein content per cell (43).

LPR normalization helps to adjust artefacts of median and other normalization approaches, which is achieved by the combined use of global- and nascent proteome analysis in the same sample. It exploits plexDIA of SILAC-labelled samples, which quantifies accurately even across abundance ratios of 100-fold (**Figure 2, Supplementary Figures 1 and 2**). This contrasts with label-free benchmarks that use multi-species mixtures to evaluate quantification, however usually only up to 10-fold differences (44). Conceptually, LPR normalization is similar to a method for normalizing phosphoproteomic data when protein phosphorylation is globally inhibited, which enabled quantification of extreme phosphorylation regulation (45). That approach, however, used ≤50 quantifiable phosphopeptides, whereas our 4–6 h pulse labelling yields hundreds to thousands of quantified NSPs. The directLFQ algorithm scales data using selected protein subsets, such as contaminants and housekeeping proteins (11), but these are chosen on the assumption that their abundance is constant — an assumption that is neither tested before nor after correction. Deriving the scaling factor from the median ratio of all NSPs quantified across samples, as we do here, is less biased.

Although eIF4A1 and eIF4A2 share ∼90% sequence identity, only eIF4A1 is essential for cap-dependent translation initiation (31, 46). Consistent with these reports, eIF4A1 depletion strongly reduced protein synthesis, whereas eIF4A2 depletion had only minor effects on the nascent and global proteome (**Figure 5C, E; Supplementary Figure 5**). Despite this strong inhibition, not all proteins exhibited a decreased synthesis, and several were even increased upon eIF4A1 depletion (**Figure 5D**). This seems in disagreement with the canonically known function of eIF4A as a helicase that is globally required for translation initiation, however detailed studies have revealed a more complex scenario where eIF4A preferentially translates specific transcript subsets (e.g. containing G-quadruplexes in 5’UTR (37)), and where its depletion leads to enhanced translation of other subsets (38). For instance, this is the case for eIF4A2 (**Figure 5D**), an established consequence of eIF4A1 loss of function. This has been attributed to a transcriptional response (41), indicating the complexity of translational regulation even among eIFs.

Rocaglates such as silvestrol are pursued as anticancer and antiviral agents (47), and silvestrol is among the most frequently used (48). It inhibits both eIF4A1 and eIF4A2 (49) which has been attributed to increased eIF4A–RNA binding affinity (37). In this light, our observation that treatment with silvestrol resulted in proteome changes that differed both from eIF4A1 and eIF4A2 depletion seems paradoxical. However this phenomenon has been observed before, revealing that silvestrol remodels translation in a more complex manner than only inhibiting eIF4A, and that it can promote rather than inhibit translation by engaging with other translation factors including eEF1ε1 (38). This introduces secondary effects not seen after clean knock-out of eIF4A1 or eIF4A2, thus explaining how non-overlapping proteome responses emerge.

In conclusion, LPR normalization combines nascent- and global proteome analysis to measure global changes in protein synthesis, and with SILAC plexDIA quantifies accurately up to 100-fold differences between labels. Applying the workflow exposed marked differences between the effects of depleting eIF4A1 or eIF4A2 and of inhibiting both with silvestrol. We expect the approach to prove useful for studying cellular stress responses, testing therapeutic compounds, and investigating regulators of mRNA translation.

## Methods

### Simulation of nascent proteome data

To illustrate the effect of different normalization strategies on nascent proteome data with a known ground truth, we simulated a pulsed SILAC experiment comparing control and treated cells, in which the treatment globally represses protein synthesis. All simulations were performed in R (version 4.3.3) with a fixed random seed to ensure reproducibility. A total of 3,000 quantified NSPs were simulated, each randomly assigned to one of three groups. The majority of proteins (88%) followed the global inhibition, with true log2 fold changes (treated/control) drawn from a normal distribution with a mean of −1.07 and a standard deviation (SD) of 0.30. A smaller group (8%) was largely unaffected by the treatment (mean −0.15, SD 0.25), and 4% of proteins were induced (mean 0.90, SD 0.30). Based on its true fold change, each protein was classified as truly down-regulated (log2 fold change < −0.585), truly up-regulated (> 0.585) or unchanged. Measured log2 SILAC ratios of the enriched NSPs were simulated for four replicates as the true fold change plus a systematic mixing error and protein-specific measurement noise. The mixing error, representing unequal mixing of control and treated lysates before NSP enrichment, was drawn per replicate from a normal distribution (mean 0.50, SD 0.12 log2 units; about 1.4-fold). The measurement noise depended on protein intensity, with an SD ranging from 0.18 to 0.43 log2 units. The global proteome before enrichment was simulated as 900 log2 ratios of labelled over unlabelled protein per condition and replicate (SD 0.55), with the control distribution (H/L) centred at −2.67 and the treated distribution (M/L) shifted by the median true fold change of the NSPs. The LPR factor of each replicate was calculated as the difference between the median log2 M/L and H/L ratios.

The simulated NSP ratios were either left unnormalized, centred on the median of each replicate (standard median normalization), or median-centred and then shifted by the LPR factor of the same replicate (LPR normalization). Differential abundance was assessed using a moderated one-sample t-test with empirical Bayes variance shrinkage and Benjamini-Hochberg correction. Proteins with an adjusted p-value below 0.05 and an absolute log2 fold change above 0.585 were considered regulated, and each call was classified against the ground truth as true or false positive, or true or false negative.

### Cell culture

HeLa cells, (ATCC cat. no. CCL-2), were grown in DMEM high glucose medium (Gibco) supplemented with 2 mM L-glutamine, 10 % (v/v) fetal bovine serum (FBS) (Gibco) and additional 2 mM GlutaMAX (Gibco). RKO cells (ATCC cat. no. CRL-2577) were grown in RPMI 1640 medium (Gibco) supplemented with 10% FBS and 4 mM GlutaMAX (Gibco).

Genetically modified RKO cells were prepared as described previously (50). Specifically, a clonal cell line of RKO colorectal cancer cells, with inducible spCas9 and eGFP expression, served as template for the generation of a cell line with inducible eIF depletion. The RKO cell lines were further modified through lentiviral transduction, introducing a vector containing a neomycin resistance gene, Thy1.1 selection marker and two sgRNAs targeting the EIF4A1 or EIF4A2 gene. Modified RKO cell lines were selected through antibiotic selection with geneticin (G418) and fluorescence activated cell sorting (FACS) selecting Thy1.1-positive cells. Doxycycline (200 ng/mL) was added to induce the expression of spCas9. In order to control for the effects of doxycycline treatment and sgRNA transduction and expression, RKO cells expressing sgRNA targeting the integration site of Adeno-associated virus DNA on chromosome 19 (q13.3-qter, AAV1) (51) were included as controls for all experiments.

For the preparation of the SILAC dilution benchmark samples, HeLa cells were grown in 10 cm dishes with high glucose DMEM, supplemented with dialyzed fetal bovine serum (Gibco) and heavy-(^13^C_6_^15^N_4_-Arg, ^13^C_6_^15^N_2_-Lys), medium-(^13^C_6_-Arg, D_4_-Lys) or light isotope-containing lysine and arginine for 10 days. For the pSILAC/AHA mixing benchmark, HeLa cells were seeded in 15 cm dishes in high glucose DMEM supplemented with 10% (v/v) FBS and 1% (v/v) penicillin-streptomycin (Gibco) one day before metabolic labelling. In all other experiments, cells were grown in 6-well plates.

### Cell viability assay

Cell viability assays were performed using the CellTiter Glo-2.0 kit (Promega), according to the manufactureŕs instructions. One thousand RKO cells per well were seeded in white 96-well plates (Nunclon, Thermo Fisher Scientific). Cellular ATP-content was measured 0 h, 24 h, 48 h and 72 h after seeding using a 96-well luminescence plate reader (Mithras LB 940, Berthold Technologies).

### Metabolic labelling of newly synthesized proteins

Nascent proteome analysis was performed essentially as described before (7, 52). Cells were first washed with warm PBS and incubated with DMEM high glucose- or RPMI 1640 medium deprived of methionine, arginine and lysine for 45 min. The pulsed SILAC and AHA labelling was carried out with methionine-free DMEM high glucose- or RPMI 1640 medium containing heavy-(^13^C_6_^15^N_4_-Arg, ^13^C_6_^15^N_2_-Lys) or medium (^13^C_6_-Arg, D_4_-Lys) lysine, arginine and 100 µM AHA for 4 h for HeLa cells or 6 h for RKO cells.

### Cell lysis and ultrasound sonication

Cells were lysed with lysis buffer containing 1% Sodium-dodecylsulfate (SDS), 300 mM HEPES (pH 8.0) and cOmplete EDTA-free protease inhibitor cocktail (Merck). Cell lysates were sonicated in AFA-tube TPX strips (Covaris), using a Covaris LE220R-Plus for 300 s at 325 peak power with a duty factor of 50 %, 200 cycles per burst, average power of 162.5. The dithering parameters were set to ± 5 mm in x and z direction and 4.5 mm in y direction at a speed of 20 mm/s. Protein concentrations of the sonicated lysates were determined using a BCA assay (Pierce). A total of 100 µg (50 µg per condition) protein was used as input for the enrichments.

### Semi-automated enrichment of newly synthesized proteins

A semi-automated method was used for the enrichment of newly synthesized proteins as described in (9). Combined lysates (100 µg total protein input) were diluted to a total volume 150 µL using lysis buffer. In order to prevent the coupling of proteins containing strongly nucleophilic cysteine to the beads (53), the samples were alkylated by addition 3.4 µL of 600 mM iodoacetamide (IAA) for 20 min at room temperature. Subsequently, 20 µL of magnetic alkyne agarose (MAA) beads (20 % suspension) prepared as described (9), diluted in lysis buffer and 9.3 µL of [3+2] cyclo addition reaction mixture, containing 21.62 mM CuSO_4_, 108.11 mM Tris-hydroxypropyltriazolylmethylamine (THPTA), 216.22 mM pimagedine hydrochloride and 216.22 mM sodium ascorbate. Next, the plate was removed from the Bravo platform, sealed using VersiCap Mat 96-well flat cap strips (Thermo Fisher Scientific) and incubated in a thermal shaker for 2 h at 40 °C. Following the coupling of the AHA-containing NSPs, the plate was unsealed and moved back onto the orbital shaker on the Bravo platform, and the supernatant was removed, by placing the sample plate on a magnetic rack (ALPAQUA MAGNUM FLX enhanced universal magnet) for 30 s and aspirating the supernatant in two steps using tips from position 6 and dispensed in the waste plate. The beads were subsequently washed with 150 µL milliQ water. After addition of the liquid to the beads on the orbital shaker, the plate was moved to the heating station where the beads were mixed by pipetting up and down 8 times with a constant flow rate of 300 µL/s, to prevent aggregation of the beads. Next, the plate was transferred onto the magnetic rack and the supernatant was removed. The bead-bound NSPs were subsequently reduced and alkylated by addition of 150 µL of 10 mM Tris(2-carboxylethyl)phosphine (TCEP) and 40 mM 2-chloroacetaminde (CAA), dissolved in 100 mM Tris-HCl buffer (pH 8.0), containing 200 mM NaCl, 0.8 mM Ethylendiamintetraacetic acid (EDTA), 0.8% SDS, followed by incubation on the heating station at 70 °C for 20 min and subsequently at 20 °C for 15 min on the orbital shaker. Next, the beads were washed three times with 1% SDS dissolved in 100 mM Tris-HCl (pH 8.0), 250 mM NaCl and 1 mM EDTA buffer, once with milliQ H_2_O, three times with 6 M Guanidine-HCl in 100 mM Tris-HCl (pH 8.0) and three times with 70% ethanol, in consecutive washing steps of 150 µL each. Following the washing steps, the beads were resuspended in 50 µL 100 mM Ammonium bicarbonate buffer (pH 8.0). Proteins were digested off the beads by adding 6 µL of 0.1 µg/µL sequencing grade Trypsin (Promega), diluted in 50 mM acetic acid, for 16 h at 37 °C, which was performed in a thermal shaker after sealing the plate with VersiCap Mat 96-well flat cap strips.

### pSILAC/AHA mixing benchmark: sample preparation and data acquisition

For the pSILAC/AHA mixing benchmark (SILAC benchmark II), previously published (semi-)automated workflows for global proteome analysis (autoSP3) (56) and for pulsed SILAC and AHA labelling with enrichment of newly synthesized proteins (7, 9) were used with minor adaptations. HeLa cells were depleted of methionine, arginine and lysine for 45 min in SILAC DMEM (AthenaES) lacking these amino acids, supplemented with 10% (v/v) dialyzed FBS (Gibco), 1 mM sodium pyruvate (Gibco), 1× GlutaMAX (Gibco), 3.7 g/L sodium bicarbonate (Sigma Aldrich), 0.105 mg/mL L-leucine and 0.2 mg/mL L-proline (Sigma Aldrich) and 1% (v/v) penicillin-streptomycin (Gibco), pH 7.4. Cells were then pulse-labelled for 4 h in the same medium containing 0.2 mg/mL AHA and either heavy (¹³C₆¹⁵N₄-Arg, ¹³C₆¹⁵N₂-Lys) or medium (¹³C₆-Arg, D₄-Lys) SILAC amino acids (0.084 mg/mL arginine, 0.146 mg/mL lysine). Cells were washed twice with PBS, harvested by scraping, pelleted (1,000 × g, 10 min, 4 °C) and stored at −80 °C.

Cell pellets were lysed in 200 µL lysis buffer (1% SDS, 300 mM HEPES pH 8.0, 1× cOmplete protease inhibitor cocktail (Roche)), and lysates of the same SILAC label were pooled. Lysates were sonicated on ice (Branson sonicator, 10% amplitude, 10 s on/10 s off for 4.5 min) and cleared by centrifugation (16,000 × g, 30 min, 12 °C). Protein concentrations were determined by BCA assay (Pierce, Thermo Scientific). For global proteome analysis, 50 µg protein per SILAC label was processed using the autoSP3 workflow on a Bravo liquid handling platform (Agilent) as described above, except that proteins were digested in 40 µL of 50 mM triethylammonium bicarbonate (TEAB) with sequencing-grade trypsin (1:50 enzyme to protein) for 16 h at 37 °C. Digests were acidified with 10% TFA, and heavy- and medium-labelled peptides were mixed at H:M ratios of 0:100, 10:90, 30:70, 50:50, 70:30, 90:10 and 100:0.

For nascent proteome analysis, heavy- and medium-labelled lysates were mixed at the same H:M ratios to a total of 100 µg protein, adjusted to 120 µL with lysis buffer, and NSPs were enriched as described above, except that 135 µL was used for all washing steps after click-chemistry coupling. On-bead digestion was performed in 50 µL of 50 mM TEAB with 0.5 µg sequencing-grade trypsin for 16 h at 37 °C. Peptides were separated from the beads, lyophilised and purified using the autoSP3 peptide clean-up (9), and resuspended in 15 µL 0.1% formic acid. Global proteome samples were analysed in (n = 6) and NSP samples in (n = 3) replicates per mix.

### Automated peptide purification

The automated SP3 protocol was used for the purification of peptides by processing of 16-96 samples in parallel on a Bravo liquid handling robot (54). Following protein digestion, the peptide-containing supernatant was transferred from the sample plate on position 7 onto a new plate on position 8, and peptides were lyophilized using a UNIVAPO-150H vacuum concentrator, coupled to a UNICRYO MC2 cooling trap and UNITHERM 4/14 D closed circuit cooler (UNIEQUIP). On the Bravo platform, magnetic carboxylate Sera-Mag Speed Beads (Fisher Scientific) were diluted to 100 µg/µL in 10 % formic acid and 5 µL were added to each lyophilized sample. Aggregation of the peptides onto beads was induced via addition of 195 µL acetonitrile and incubation for 18 min, while shaking at 100 rpm on the orbital shaker. Next, the supernatant was removed from the magnetic rack in two steps. The beads were washed 2 times with 180 µL acetonitrile and subsequently dried. In the final steps the beads were resuspended in 20 µL 0.1 % formic acid in water and sonicated in an Ultrasonic Cleaner USC-T (VWR) for 10 min and the supernatant was transferred to a new plate. The purified peptides were dissolved in 0.1 % formic acid and used for LC-MS/MS analysis.

### Preparation of proteomics samples without enrichment of newly synthesized proteins

20 µg of either pulse- or stable SILAC labelled cell lysate were used for the preparation of non-enriched proteome samples. Lysates were obtained from the same samples, that were used for the enrichment of newly synthesized proteins. Protein concentration of the lysates was determined using a BCA assay (Pierce). Proteome samples were prepared using the automated SP3 protocol (54) as described above.

### LC-MS/MS

MS-based proteome analysis was performed using an EASY-nLC 1200 system (Thermo Fisher Scientific) coupled to a Q Exactive HF mass spectrometer (Thermo Fisher Scientific). Peptides were separated by reverse-phase liquid chromatography using 0.1 % formic acid (solvent A) and 80 % acetonitrile (solvent B) as mobile phases. Peptide separation occurred on an Acclaim PepMap trap column (Thermo Fisher Scientific, C18, 20 mm × 100 μm, 5 μm C18 particles, 100 Å pore size) and a nanoEase M/Z peptide BEH C18 analytical column (Waters, 250 mm × 75 μm 1/PK, 130 Å, 1.7 μm). The samples were loaded onto the trap column with constant flow of solvent A at a maximum pressure of 800 bar. The analytical column was equilibrated with 2 μL solvent A at a maximum pressure of 600 bar heated to 55 °C using a HotSleeve+ column oven (Analytical SALES & SERVICES). Peptides were eluted with a constant flow rate of 300 nL/min. Concentration of solvent B was gradually increased during the elution of the peptides in either of two different HPLC gradients used in this study. The gradient used for the analysis of the newly synthesized proteome samples and proteome samples of HeLa cells started with 4% solvent B and was increased to 6% in the first 1 min, to 27% until 51 min, and to 44% until 70 min. This was followed by a rinsing and re-equilibration step at 95% and 5% solvent B, respectively, each for 10 min. The gradient used for the analysis of non-enriched (global proteome) samples started at 3% solvent B, which was increased linearly to 8% B at 4 min, 10% B at 6 min, 32% B at 68 min and 50% B at 86 min. The column was then washed with 100% B from 87 to 94 min and re-equilibrated with 3% B for 10 min from 95 min. Eluting peptides were ionized and injected into the mass spectrometer, using the Nanospray flex ion source (Thermo Fisher Scientific) and a Sharp Singularity nESI emitter (ID = 20 µm, OD = 365 µm, L = 7 cm, α = 7.5°) (FOSSILIONTECH), connected to a SIMPLE LINK UNO-32 (FOSSILIONTECH). A static spray voltage of 2.5 kV was applied to the emitter and the capillary temperature of the ion transfer tube was set to 275 °C.

The **Q Exactive HF mass spectrometer** was operated in the data-dependent (DDA) or data-independent (DIA) mode for the three different acquisition methods evaluated in this study. For DDA measurements of CHX-treated HeLa cells, a full scan range of 375-1500 m/z, Orbitrap resolution of 60000 FWHM, automatic gain control (AGC) target of 3e6 and maximum injection time of 32 ms was set. Data-dependent MSMS spectra were acquired using a Top 20 scheme, using a fixed scan range from 200-2000 m/z and fixed first mass of 110 m/z. The quadrupole isolation window was set to 2.0 m/z and normalized collision energy was set to 26. The Orbitrap resolution was set to 15000 FWHM with an AGC target of 1e5 and maximum injection time of 50 ms. MSMS spectra were acquired in profile mode and a charge state exclusion of 1, 5-8 & >8 was defined. An intensity threshold of 2e4 and a minimum AGC target of 1e3 was set.

Two different DIA methods were used. For the analysis of the global RKO cell proteome (without NSP enrichment) the method used a full scan range of 400-1000 m/z with Orbitrap resolution of 60000 FWHM, 3e6 AGC target and maximum injection time of 20 ms. Data-independent MSMS spectra were acquired with an Orbitrap resolution of 30000 FWHM, maximum injection time of 50 ms and AGC target of 1e6, using 26 equally sized, 1 Th overlapping isolation windows with a width of 23.3 m/z. The normalized collision energy for the fragmentation of precursor ions was set to 27 and a fixed first mass of 200 m/z was set for the acquisition of the MSMS spectra. The normalized collision energy for the fragmentation of precursor ions was set to 27 and a fixed first mass of 200 m/z was set for the acquisition of the MSMS spectra. For the analysis of all other samples, the DIA method used a full scan range of 400-1000 m/z, Orbitrap resolution of 60000 FWHM, 3e6 AGC target and maximum injection time of 40 ms. Data-independent MSMS spectra were acquired with an Orbitrap resolution of 30000 FWHM, maximum injection time of 40 ms and AGC target of 1e6, using 28 equally sized, 1 Th overlapping isolation windows with a width of 22.0 m/z. The normalized collision energy for the fragmentation of precursor ions was set to 27 and a fixed first mass of 200 m/z was set for the acquisition of the MSMS spectra.

Samples of the pSILAC/AHA mixing benchmark were analysed on an EASY-nLC 1200 system coupled to an **Orbitrap Fusion mass spectrometer** (both Thermo Fisher Scientific). Peptides were separated at a flow rate of 300 nL/min using solvent A (0.1% formic acid in water) and solvent B (0.1% formic acid in 80% acetonitrile). The gradient started at 2% B for 3 min, increased to 38% B over 100 min and to 52% B over 10 min, followed by a wash at 100% B for 6 min and re-equilibration at 2% B for 11 min (total method duration 130 min). Peptides were ionized by nanoelectrospray at a static spray voltage of 2.5 kV, with the ion transfer tube at 275 °C, and internal mass calibration was performed using a lock mass of m/z 445.12003. Full MS spectra were acquired in the Orbitrap over m/z 350–1000 at a resolution of 60,000, with a standard AGC target, a maximum injection time of 60 ms and an RF lens setting of 60%. DIA MS/MS spectra were acquired using 44 non-overlapping isolation windows of variable width (9.5–23.5 m/z) covering m/z 407–961. Precursors were fragmented by HCD at a normalized collision energy of 33%, and fragment spectra were recorded in the Orbitrap at a resolution of 30,000, with a first mass of m/z 145, a normalized AGC target of 800%, a maximum injection time of 60 ms and an RF lens setting of 60%. Data were processed with DIA-NN (plexDIA) as described below.

### Proteomic data analysis

Raw files from DDA measurements were processed using Maxquant (version 2.0.3) and the Andromeda search engine (55), using a human proteome fasta file, retrieved from the SwissProt database (version from February 2021 with 20934 entries). The enzymatic digestion was set to Trypsin/P and a maximum of 2 missed cleavages per peptide were allowed. For the analysis of NSP data, raw files of both the NSP and global proteome samples were processed together, using Maxquant (version 2.0.3). The multiplicity was set to 3, comprising of a light channel, an medium channel with Arg6 and Lys4 and heavy channel with Arg10 and Lys8. Cysteine carbamidomethylation was set as fixed modification, whereas methionine oxidation, N-terminal acetylation, and deamidation of Asparagine and Glutamine were set as variable peptide modifications. Samples were processed with the match-between-runs disabled, and the re-quantify function either disabled or enabled. Default settings were used for other search functions. Minimum peptide length was set to 7 and max peptide mass was set to 4600 Da. PSM-, Protein- and site decoy fraction FDR were set to 1 %. The minimum delta score threshold for unmodified peptides was set to 6, and 40 for modified peptides. Only for the analysis of the SILAC labelled Hela benchmark samples, the match-between-runs function was enabled. Unique and razor peptides were used for quantification and normalized SILAC ratios and iBAQ values were calculated. The minimum ratio count was set to 0 to not exclude identifications in single SILAC channels.

Raw files from DIA measurements were analysed using DIA-NN (56) (version 1.8.1). A predicted spectral library was generated from the fasta file that was also used in the MaxQuant searches. Additionally, a FASTA file containing common protein contaminants was added for the spectral library prediction (57). Default settings were used for the spectral library prediction, with the addition of methionine oxidation as variable modification. For the processing of the raw files, the default settings of DIA-NN were used with additional functions from the plexDIA module enabled (8). Three SILAC channels with mass shifts corresponding to Lys, Lys4 (+4.025107 Da), Lys8 (+8.014199 Da), Arg, Arg6 (+6.020129 Da), Arg10 (+10.008269 Da) and an additional decoy channel with lysine (+12.0033 Da) and arginine (+13.9964 Da) were registered. Translation of retention times between peptides within the same elution group was enabled. The first ^13^C-isotopic peak and monoisotopic peak was included for the quantification and the MS1 deconvolution level was set to 2. Peptide length range was set from 7-30, precursor charge rate was set from 1-4, mass of charge (m/z) range of the precursors was set from 300-1800 and fragment ion m/z range was set from 200-1800. Precursor FDR was set to 1 %. Precursor matrix output tables were filtered for FDR < 0.01 and additionally for channel q-value < 0.01 and translated q-value < 0.01. The MBR function in DIA-NN was disabled.

### Experimental design and statistical rationale

For nascent proteome analysis we used a method combining pulse-labeling of cells with AHA and SILAC-labeled amino acids for the enrichment and quantification of newly synthesized proteins (7, 9, 52). To determine how protein synthesis depends on cellular perturbations, we treated cells with CHX or silvestrol or performed inducible knock-out of eIF4A1 and eIF4A2 under conditions as specified above. Pulse-labelling with SILAC and AHA was performed for a defined short time period (4 or 6 h) to measure direct effects of these perturbations. Proteome analysis was performed with and without enrichment to determine proteome effects in the nascent and global proteome, respectively, and to derive the LPR normalization factor.

#### Benchmark experiments

The SILAC dilution benchmark consisted of six mixtures of light-, medium- and heavy-labelled HeLa lysates (33% to 1% labelled protein per label) and a light-only sample, each analysed in two technical replicates. Linearity was assessed by linear regression of measured versus expected median log2 ratios, precision by the coefficient of variation (CV) of ratios between replicates, and accuracy by the deviation of measured from expected ratios. In the pSILAC/AHA mixing benchmark, heavy- and medium-labelled lysates from independently cultured plates were mixed at seven defined H:M ratios; each mix was analysed in six (global proteome) and three (NSP enrichment) biological replicates, each derived from a separate pair of cultures. Because the true H/M ratio of each mix is known, normalization strategies were evaluated by the difference between the median normalized log2 H/M ratio and the expected value. *Perturbation experiments.* The CHX experiment comprised DMSO-treated control cells and cells treated with 0.1, 1, 10 or 100 µg/mL CHX, in three biological replicates per condition. For eIF4A1 and eIF4A2 depletion, cells expressing an sgRNA targeting the AAV1 safe-harbour locus served as controls, and DMSO-treated cells as controls for silvestrol treatment, with 3 biological replicates per condition. Each replicate was analysed both before NSP enrichment (global proteome, LPR factor) and after enrichment (nascent proteome). The LPR factor was calculated per experiment as the difference between the mean of the replicate medians of the log2 labelled/unlabelled ratios of treated and control cells.

### Statistical analysis

Only unique protein groups with SILAC ratios in at least two of three replicates were included in differential analysis. Differences between conditions were tested with an empirical Bayes moderated t-test using limma and DEqMS (58, 59), with the number of precursors per protein group included in the variance estimation. P-values were adjusted for multiple testing using the Benjamini-Hochberg method, and proteins were considered regulated at |log2 fold change| > 0.585 (1.5-fold), adjusted p < 0.05 and a CV < 20% across replicates. For LPR-normalized data, fold changes were shifted by the LPR factor, and moderated t-statistics and p-values were recalculated using the standard errors and degrees of freedom of the median-normalized analysis. Unnormalized fold changes were obtained by adding the median offset between raw per-protein log2 ratios and median-normalized fold changes. Agreement between conditions was assessed by Pearson correlation of log2 fold changes.

After mass spectrometric analysis and data processing, the output tables from MaxQuant (“ProteinGroups.txt”, “evidence.txt”) and DIA-NN (“report.pr_matrix_channels_translated.tsv” were processed in the R software environment (version 4.0.3) using custom scripts. Identified contaminants were removed and protein abundance was calculated using the MaxLFQ algorithm, applied to the individual SILAC channels, using the iq (version 1.9.6) R package function “process_long_format()” (60). MS2-based quantification, more specifically the “precursor.translated” metric was used for the MaxLFQ protein intensity normalization. Protein-group SILAC ratios were calculated for each sample using the calculated LFQ values. Intensity-based absolute quantification (iBAQ) normalization for individual proteins was carried out by dividing the “PG.Quantity” metric (for proteome data), or the summed “precursor.translated” metric (for nascent proteome data), by the number tryptic peptides (length ≤ 30 aa & ≥ 7 aa). For differential expression analysis, only unique protein groups (single Uniprot identifier) with a minimum of 2 SILAC ratios values in 3 replicates were used. Differential expression tests were carried out using the Limma (version 3.46.0) (58) and DEqMS (version 1.8.0) (59) R/Bioconductor packages, by fitting the data onto a linear model and performing an empirical Bayes moderated t-test. Additional normalization of the global proteome data was carried out using variance stabilization normalization via the “normalizeVSN” function of the limma R/Bioconductor package. The number of precursors of each protein group, with consideration of modified peptide sequences and charge but not SILAC channels, was included as a factor for the variance estimation in DEqMS.

### Comparison of normalization strategies

To compare LPR normalization with commonly used normalization strategies, we used the NSP data of the 30:70, 50:50 and 70:30 H:M mixes of the pulsed SILAC/AHA benchmark, for which the expected log2 H/M ratios are −1.22, 0 and 1.22. Protein-level intensities of the heavy (H) and medium (M) channels were analysed separately for each mix. The analysis included only proteins quantified in both channels in all three NSP replicates. Normalization methods were applied at three levels: to the channel intensities, to the protein H/M ratios, or as a single global factor derived from the non-enriched global proteome. For intensity-level normalization, the six channel intensities of each mix were normalized as separate samples, and log2 H/M ratios were calculated afterwards for each replicate. Total intensity, mean, median, quantile, variance stabilization (VSN) and robust linear regression (RLR) normalization were performed with NormalyzerDE (version 1.31.0) (61). Cyclic loess normalization was performed with the normalizeCyclicLoess function of limma (version 3.58.1, method “fast”) (58). In addition, directLFQ (version 0.3.3) was applied to the precursor-level intensities of each mix, and the resulting protein intensities were used to calculate H/M ratios. For ratio-level normalization, log2 H/M ratios were first calculated per protein and replicate, and the three replicates were then normalized against each other. Standard median normalization centred each replicate on its median. Quantile normalization was performed with preprocessCore (version 1.64.0) (62), and cyclic loess normalization with limma as above. As a reference, the ratios were also analysed without any normalization.

For the global-factor methods, the NSP ratios of each replicate were first median-centred and then shifted by a single factor derived from the non-enriched global proteome of the same mix. For LPR normalization, this factor was the median log2 H/M ratio across all proteins quantified in both channels in the global proteome. Because H- and M-labelled lysates were already combined in these samples, the light, pre-existing proteome cancels, and the median H/M ratio corresponds to the difference between the median H/L and M/L ratios. For the channel-sum ratio, the factor was calculated as the log2 ratio of the summed H over the summed M intensities of all quantified proteins in each global proteome replicate, and the median across replicates was used. For every strategy, the mean log2 H/M ratio of each protein across the three replicates was estimated with a linear model and empirical Bayes moderation.

### Pathway enrichment analysis

To identify the biological processes affected by the choice of normalization, NSPs from the CHX (0.1 µg/mL) and eIF4A1 depletion experiments were grouped according to their regulation after standard median and after LPR normalization (log2 fold change > 0.585, adjusted p < 0.05). Groups were defined as NSPs regulated with both methods, NSPs down-regulated only with LPR normalization (hidden by median normalization), and NSPs up-regulated only with median normalization (artefacts of median normalization). Gene symbols were mapped to Entrez Gene identifiers with org.Hs.eg.db (version 3.18.0). Gene Ontology Biological Process annotations, including all parent terms, were obtained from org.Hs.eg.db and GO.db (version 3.18.0), and Reactome pathways from the Molecular Signatures Database (MSigDB, C2:CP:REACTOME, version 7.5.1). Over-representation of each term in each group was tested with a one-sided Fisher’s exact (hypergeometric) test, using all NSPs quantified in the respective experiment as background. Only terms containing 10 to 500 background proteins and at least three proteins of the tested group were considered. P-values were adjusted separately for each group and database using the Benjamini-Hochberg procedure, and terms with an adjusted p-value below 0.05 were considered enriched.

### Data availability

Protein-level abundances and fold changes for all datasets are provided in Supplementary Tables 1–4.

All mass spectrometry data have been deposited to the ProteomeXchange Consortium via the PRIDE repository (63):

SILAC dilution benchmark (Q Exactive HF): PXD050068

pSILAC/AHA mixing benchmark (Orbitrap Fusion): PXD086013

Cycloheximide dose series (plexDIA): PXD046862

Cycloheximide, DDA analysis: PXD046890

eIF4A1/2 depletion and silvestrol treatment: PXD050075

## Acknowledgements

This work was funded in part by the German Ministry of Education and Research (BMBF), as part of the National Research Node “Mass spectrometry in Systems Medicine” (MSCoreSys), under grant agreement 161L0212A.

## Author contributions

T.B., A.S.A. and J.K. conceived the study and designed the approach. T.B., L.G., E.M. and A.S.A. prepared samples and acquired data. R.K. and J.Z. generated the RKO cell lines. T.B. and A.S.A. analysed data. T.B., A.S.A. and J.K. wrote the manuscript, with input from all authors.

## Supplementary Figure Legends

**Supplementary Figure 1.**
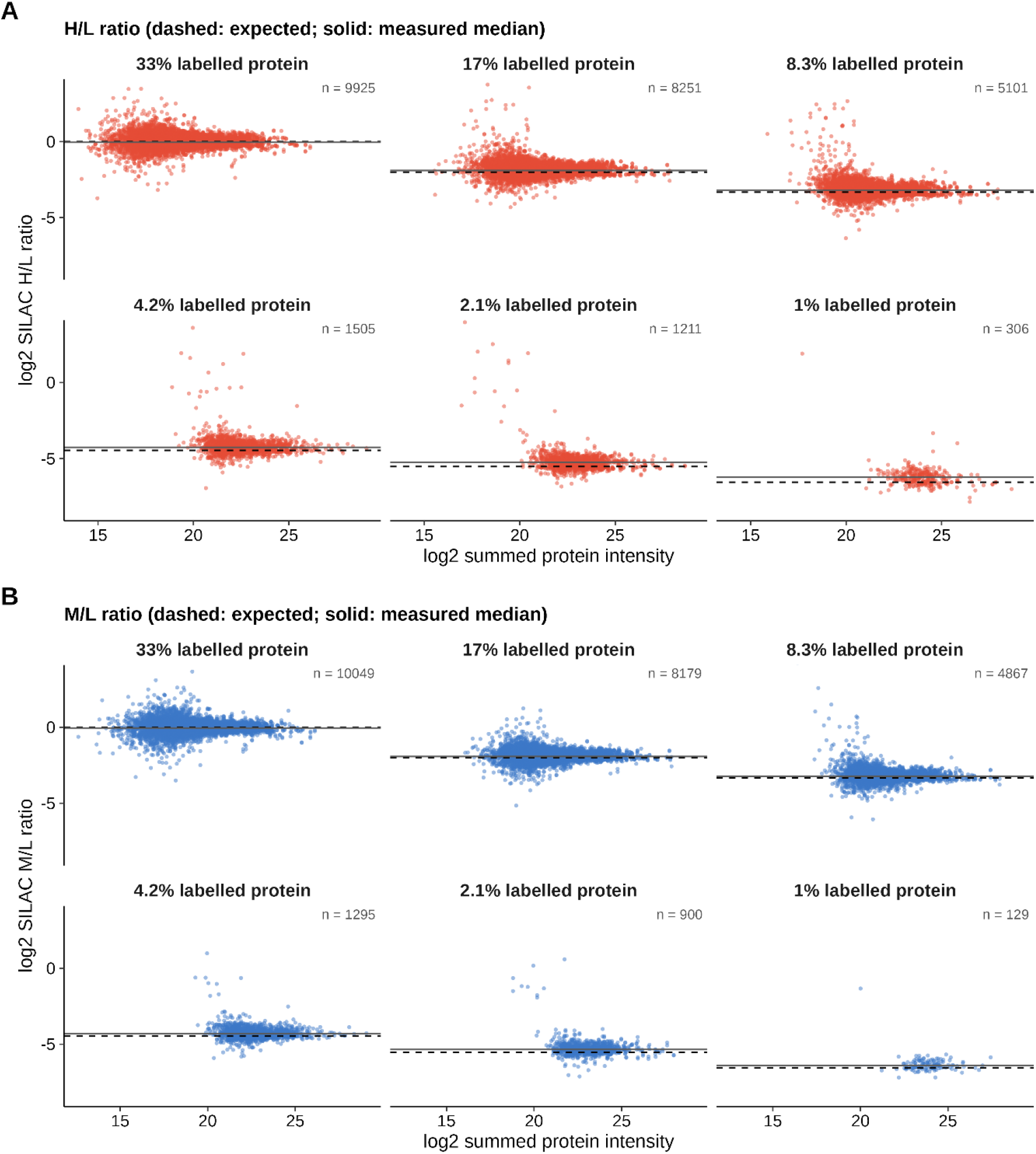
SILAC ratios are accurate across the protein intensity range. Log2 H/L (A) and M/L (B) SILAC ratios plotted against summed protein intensity for each benchmark sample of the dilution series (Figure 2A). Panels are labelled by the fraction of heavy- or medium-labelled protein (% of total protein, per label). Dashed lines indicate the expected ratio, solid lines the measured median. n, number of protein ratios from both replicates. Samples were analysed by 90 min DIA on a Q Exactive HF and processed with DIA-NN (plexDIA).

**Supplementary Figure 2.**
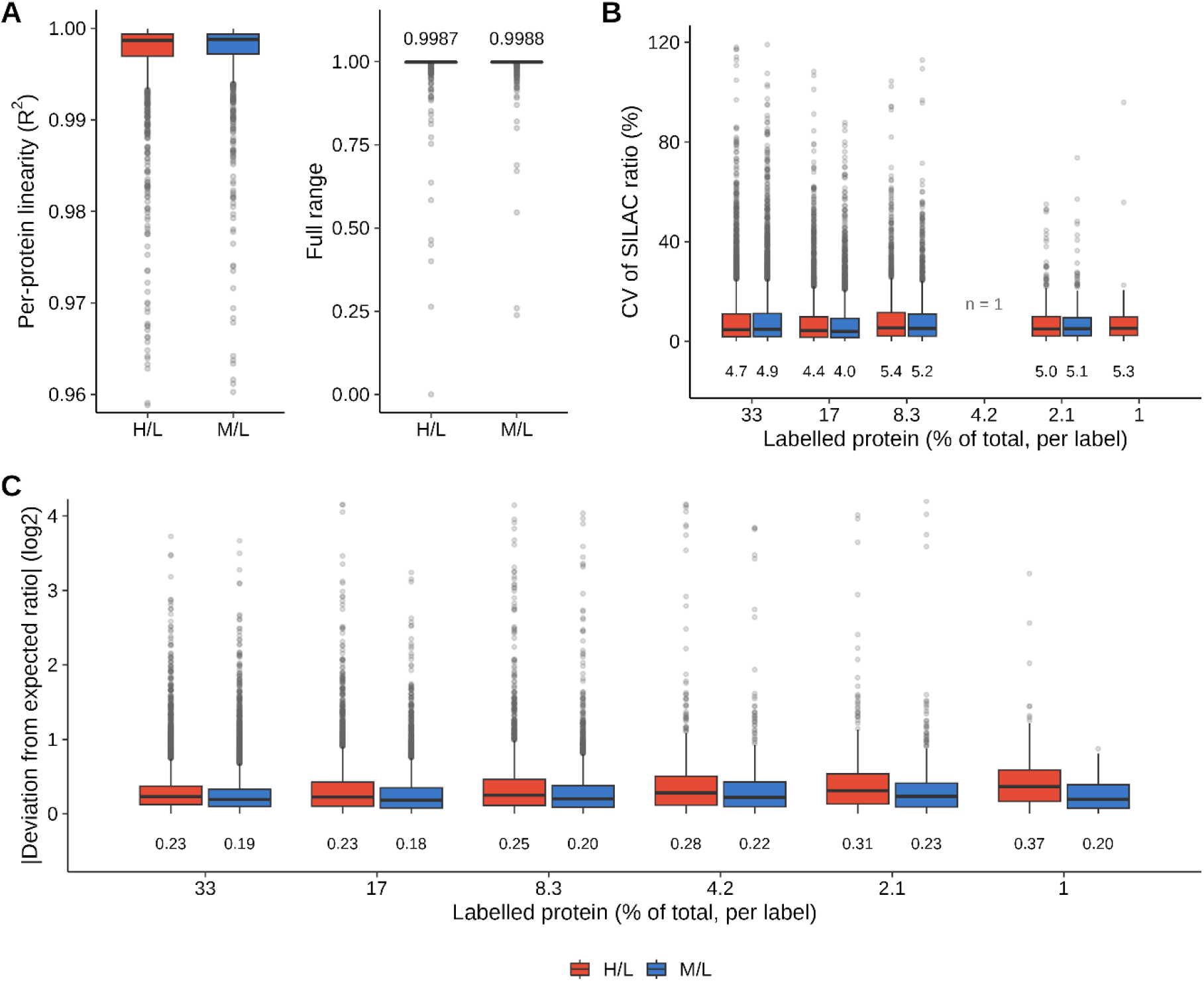
Quantitative performance of the SILAC dilution benchmark. A) Linearity of SILAC ratios per protein, shown as R² of a linear fit of measured versus expected H/L or M/L ratio for proteins quantified at ≥ 4 dilutions. The left panel is zoomed in (R² 0.96–1), the right panel shows the full range; numbers indicate median R². B) Coefficient of variation (CV) of SILAC ratios between replicates for each benchmark sample; CVs were not computed at 4.2% labelled protein (n = 1). C) Absolute deviation of measured from expected log2 SILAC ratios per protein and replicate. Numbers below the boxes indicate medians. Boxes show the interquartile range with the median, and whiskers 1.5× the interquartile range.

**Supplementary Figure 3.**
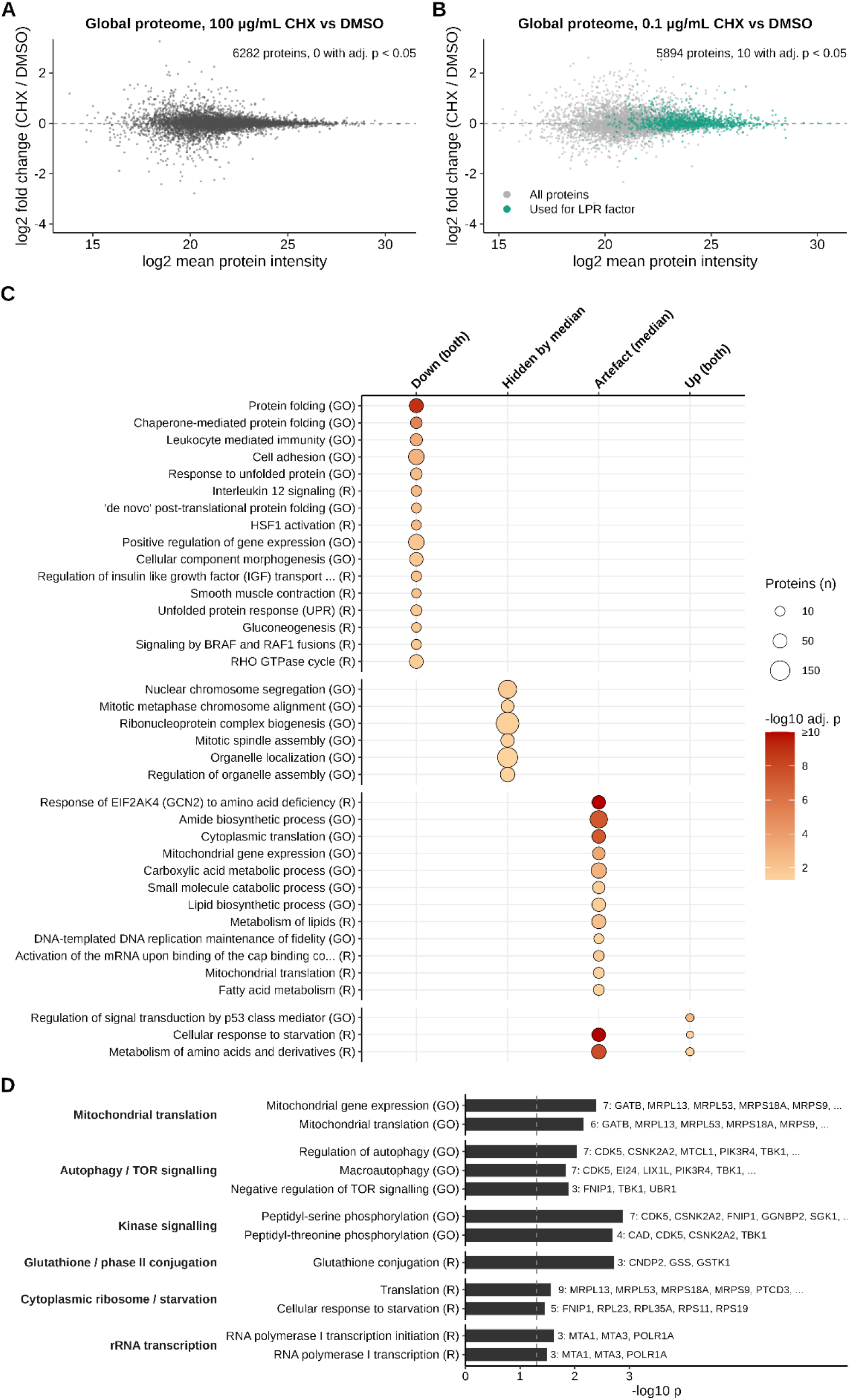
Global proteome and functional analysis of the cycloheximide experiment. A, B) Global proteome (label-free quantification, summed intensity of all SILAC channels) of HeLa cells treated for 4 h with 100 µg/mL (A) or 0.1 µg/mL (B) CHX, compared with DMSO, plotted against mean protein intensity. Numbers indicate quantified proteins and proteins with adjusted p < 0.05. In B, proteins used to calculate the LPR factor are highlighted (teal). C) GO Biological Process (GO) and Reactome (R) terms enriched among NSPs in each regulation group (down or up with both normalizations, down only with LPR, up only with median normalization). Dot size indicates the number of proteins per term and colour the adjusted p-value (one-sided Fisher’s exact test, Benjamini-Hochberg correction; background: all quantified NSPs). D) Exploratory enrichment analysis of NSPs with reversed direction (up-regulated after median normalization, down-regulated after LPR normalization), shown as unadjusted p-values; dashed line, p = 0.05. Numbers indicate the proteins per term, with example genes.

**Supplementary Figure 4.**
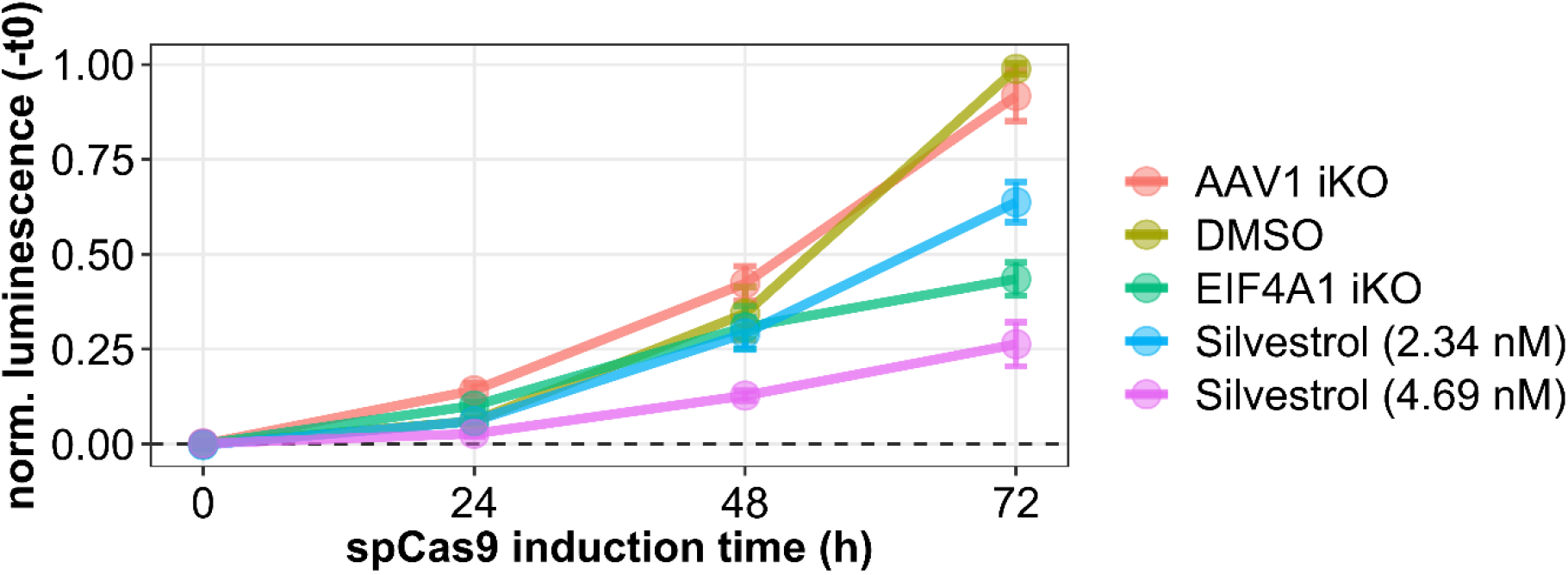
Viability of RKO cells upon eIF4A1 knockout and silvestrol treatment. Cell viability measured by CellTiter-Glo (cellular ATP content) at 0, 24, 48 and 72 h after induction of spCas9 (eIF4A1 iKO or control AAV1 iKO) or treatment with silvestrol (2.34 or 4.69 nM) or DMSO. Luminescence is normalized to t = 0.

**Supplementary Figure 5.**
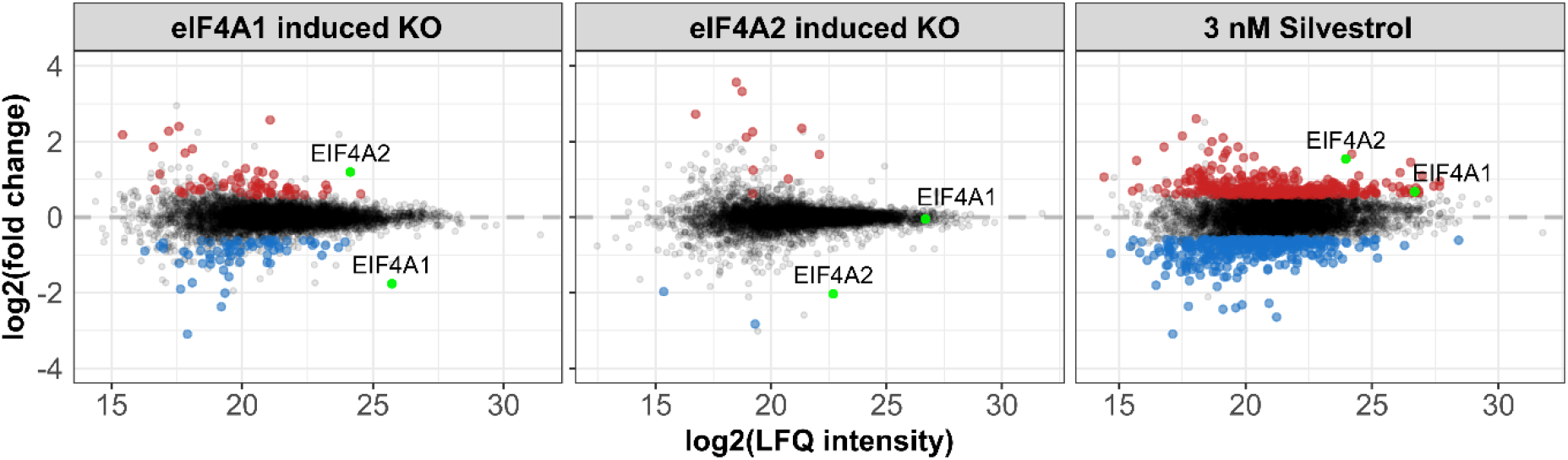
Global proteome changes upon eIF4A1 or eIF4A2 depletion and silvestrol treatment. Log2 fold changes of protein abundance (label-free quantification) after 48 h of induced eIF4A1 or eIF4A2 knockout or treatment with 3 nM silvestrol, compared with the respective controls, plotted against mean protein intensity. Significantly up- and down-regulated proteins are shown in red and blue; eIF4A1 and eIF4A2 are highlighted in green.

**Supplementary Figure 6.**
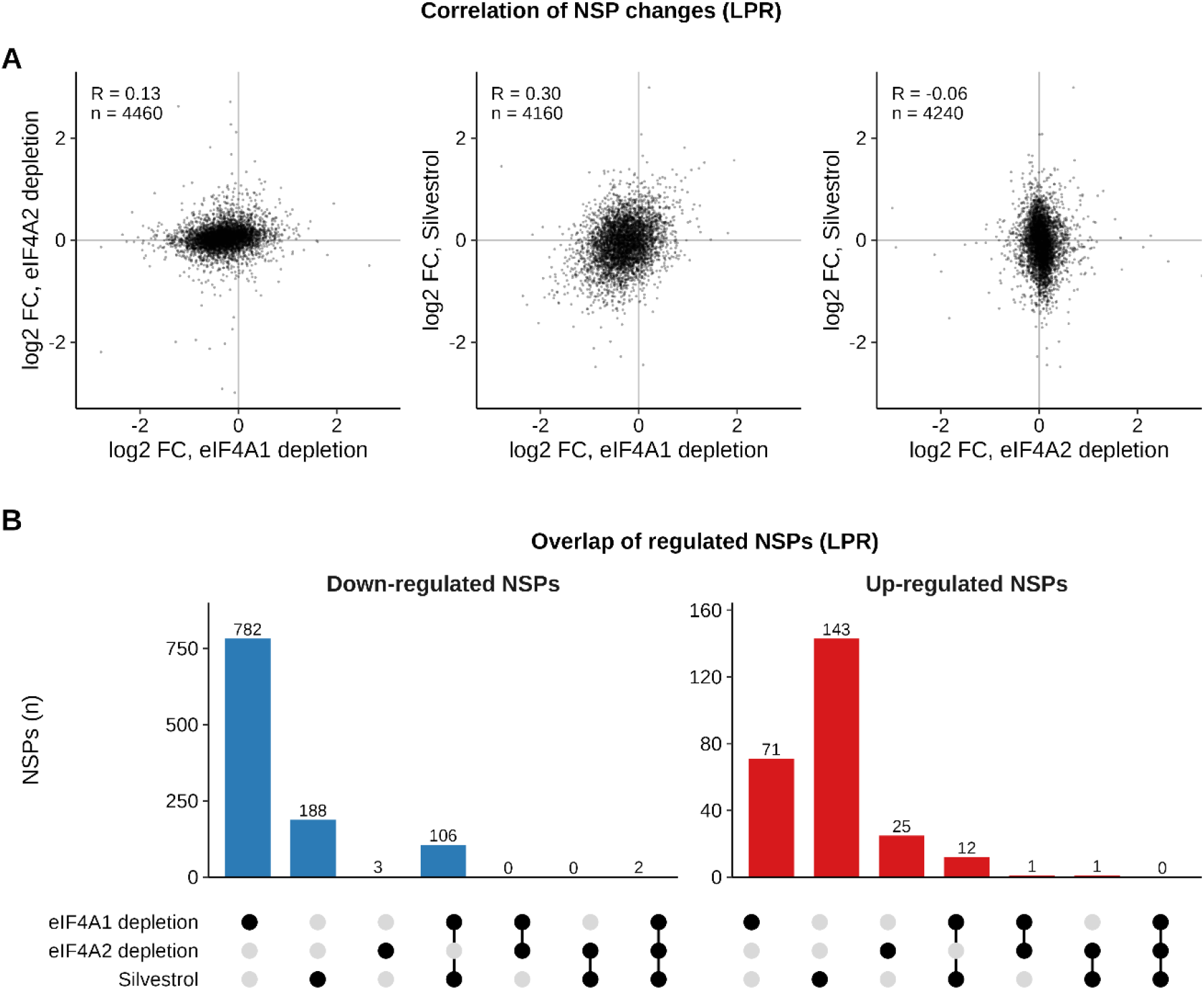
Comparison of nascent proteome changes between eIF4A1 depletion, eIF4A2 depletion and silvestrol treatment. A) Pairwise comparison of NSP log2 fold changes after LPR normalization; R, Pearson correlation coefficient; n, number of proteins quantified in both conditions. B) Overlap of down-regulated (left) and up-regulated (right) NSPs after LPR normalization (|log2 fold change| > 0.585, adjusted p < 0.05, CV < 20%). Bars show the number of proteins unique to each combination of conditions indicated below.

### Supplementary data

**Supplementary Table 1.** SILAC dilution benchmark: log2 protein intensities (sum of DIA-NN translated precursor intensities) per sample (labelled protein, % of total per label; light-only control), SILAC channel (L, M, H) and replicate.

**Supplementary Table 2.** pSILAC/AHA mixing benchmark: log2 MaxLFQ protein intensities per H:M mixing ratio, SILAC channel and replicate, for the global proteome (A) and after NSP enrichment (B).

**Supplementary Table 3.** Cycloheximide experiment: log2 protein intensities per CHX dose or DMSO control, SILAC channel and replicate, before (A) and after (B) NSP enrichment; log2 fold changes and adjusted p-values of the global proteome (100 and 0.1 µg/mL CHX vs DMSO) (C) and of NSPs (0.1 µg/mL CHX vs DMSO) without normalization, with standard median normalization and with LPR normalization (D).

**Supplementary Table 4.** eIF4A1 or eIF4A2 depletion (vs AAV1 control) and 3 nM silvestrol (vs DMSO): log2 fold changes and adjusted p-values of NSPs with standard median and LPR normalization.

